# 2,234 attempts to harmonize a multi-platform B-cell lymphoma bulk transcriptomic dataset: ComBat is superior by cross-batch prediction quality

**DOI:** 10.64898/2026.10.08.757652

**Authors:** Daniil Nikitin, Arsen Arakelyan

## Abstract

**Background:** Cross-platform transcriptomic harmonization benchmarks rely on individual success metric each: correlation with a reference profile, classifier accuracy, or a composite imported from single-cell integration. We found no published study that computes multiple orthogonal metric classes over one set of harmonization approaches and measures overlapping between the best approaches by these metrics.

**Methods:** We used the 2,234 approaches of the recent real-world benchmark on 7,174 B-cell lymphoma transcriptomes. They combine 31 harmonization methods, 14 batch-removal strategies, three imputations and two post-removal settings. We assigned its 87 polarity-adjusted scoring metrics, out of 344 registered, to five classes (global, local, distributional, structural and composite) and added three biology-facing classes: within-cohort marker correlation (L), cross-batch rank agreement of a marker panel (M), and leave-one-batch-out (LOBO) prediction of lymphoma subtypes and normal B cells (N). Each class resulted in top 112 approaches (5%), and we measured overlap by the Jaccard index.

**Results:** Harmonization methods of high harshness such as quantile normalization matched different batches to the same shape of distribution but were inferior by local and global metrics. Local and composite metrics resulted in the largest overlap of best approaches (71 shared). Classes L and M, M and N shared no approach, whereas L and N shared one approach without harmonization under the rare-batches-removed strategy, and all three overlaps were significantly below the random expectation. Class N resulted in the ComBat family in 84 of its 112 places. Class M elected only 10_mnn and 12_scanorama, methods that completely corrupted biology by correlation.

**Conclusions:** Best transcriptomic harmonization approach is highly contingent on the quality metrics used, with metric classes trade-offing between each other. We provide a six-step recipe for metric classes, their thresholds and the cross-batch tests for selecting the best approaches.

## Introduction

Technical variation between sequencing platforms, RNA extraction protocols and tissue preservation methods separates samples of the same biological class, and it distorts inference whenever it is correlated with the outcome under study (Leek et al. 2010). Algorithms that remove this variation from bulk expression matrices have been initially developed microarray integration (Lazar et al. 2013), whereas the cross-platform harmonization is more challenging, particularly, between microarrays and next-generation sequencing (NGS) (Borisov and Buzdin 2022).

Each benchmark of these algorithms declares a different criterion of success. An early 2011 empirical comparison of cross-platform normalization scored methods on inter-platform concordance of gene lists supported by ROC-like curves and bootstrap (Rudy and Valafar 2011). A later comparison scored the methods by the accuracy of a downstream subtype classifier (Thompson et al. 2016). The single-cell integration literature supplied a third criterion, a trade-off between batch effect removal and biology conservation (Luecken et al. 2021). A benchmark of batch-effect correction methods for single-cell RNA sequencing recorded the same trade-off, reporting also that quality metrics can disagree with visual inspection of the embeddings (Tran et al. 2020). Two of the metrics that this literature contributed, the k-nearest-neighbor (KNN) batch-effect test (Büttner et al. 2018) and the local inverse Simpson’s index (Korsunsky et al. 2019a), are now applied to bulk data, albeit there is still no standard of harmonization quality measurement (Borisov and Buzdin 2022).

The harmonization quality criteria are especially important in datasets where the biological contrast is small compared to the batch level technical differences. In the case of germinal center (GC) B-cell lymphomas, expression profiling splits diffuse large B-cell lymphoma (DLBCL) into germinal-center and activated B-cell types (Alizadeh et al. 2000) predicting survival after chemotherapy (Rosenwald et al. 2002). Survival in follicular lymphoma (FL) is predicted by subtle differences in immune-cell signatures of the microenvironment (Dave et al. 2004), and a 23-gene score separates FL patients into two outcome groups under immunochemotherapy (Huet et al. 2018). These small-scale transcriptomic differences are challenging to survive transfer between materials and platforms, explaining the cell-of-origin assignment reinvention for formalin-fixed paraffin-embedded (FFPE) tissue (Scott et al. 2014) and multiple cohort sources for genetic subtypes of DLBCL (Chapuy et al. 2018; Schmitz et al. 2018). The same transfer problem governs expression signatures outside lymphoma, for example microsatellite instability status (Sorokin et al. 2021), therapy prescription in colorectal (Sorokin et al. 2022) and kidney cancer (Gudkov et al. 2022) and DNA repair pathway activation in thyroid tumors (Vladimirova et al. 2021). In a recent T-cell lymphoma xenograft study, principal-component analysis (PCA) showed only a partial overlap between primary tumors and their matched xenografts, which the authors attributed to tumor content and host cells, and that variation was removed by surrogate variable analysis (SVA) (Fiore et al. 2025).

In case of the real-world batch correction scenario, the analysis is complicated by confounding between the batch variable and with the class labels (Soneson et al. 2014) which drastically reduces predictors generalizability (Parker and Leek 2012). Moreover, cross-validation could be biased under these scenarios (Varma and Simon 2006) which is amplified by selection bias among thousands of genes in transcriptomic datasets containing only tens to hundreds of samples (Ambroise and McLachlan 2002). These issues lead to prediction failures of models deployed in clinical practice (Saeb et al. 2017; Whalen et al. 2021), described as the ‘reproducibility crisis’ in the machine learning science in general (Kapoor and Narayanan 2023).

To address these current issues in transcriptomic cross-platform harmonization, we deeply analyzed the 2,234 harmonization approaches of a recently published multi-platform B-cell lymphoma benchmark (Nikitin et al. 2026 Sep 17), which calculated 344 quality metrics. The approaches combine 31 methods, 14 prior batch-removal strategies, three imputation schemes and two post-removal settings. For each metric class among local, global, distributional, composite, biomarker genes correlation before and after harmonization (class L), cross-batch agreement of biomarker genes (class M) and leave-one-batch-out (LOBO) prediction metrics (class N), we compared harmonization methods and batch removal strategies, selecting 5% best approaches by a given metric class.

The metric classes favored different harmonization methods: the global class placed 84 of its 112 best approaches under 03_limma and 13_fsmvn, whereas the local class was led by 10_mnn with 19 approaches. Class L placed 93 of its 112 best approaches under the unharmonized output 01_raw and the minimally changed 36_explobatch, so it rewarded the approaches that change expression least. Batch PCReg separated methods more strongly than batch-removal strategies (Kruskal-Wallis effect size ε² = 0.739 against 0.100), with the RNA-seq-only strategy highest on batch PCReg and the FFPE-only strategy least often passing the joint class L and M quality check. Local and composite metrics resulted in the largest overlap (Jaccard index 0.464, 71 shared approaches). Classes L and M, M and N - shared no approach, whereas L and N shared one approach, the unharmonized output 01_raw under the rare-batches-removed strategy with softimpute imputation and post-removal. All the three overlaps between L, M and N best approaches were significantly below the random exprectation (raw hypergeometric p ≤ 0.020). The 15 approaches selected by the published clustermap shared none of the 112 that class N elected. Class N resulted in the ComBat family (05_combat, 29_combat_ref, 06_combat_seq and 08_inmoose_combatseq) in 84 of its 112 places. Class M elected two methods, 10_mnn and 12_scanorama, whose median marker correlation was −0.001 and −0.008. The four ComBat family methods demonstrated best generalizability while preserving biomarkers correlation, in contrast with feature specific quantile normalization (FSQN) and surrogate variable analysis (SVA) methods shown as top performing ones in the previous benchmark.

Finally, we propose a novel six step framework for harmonization quality metrics usage to select best harmonization approaches in other bulk transcriptomic datasets, including the non-B cell lymphoma ones. The framework and the metrics can be implemented and calculated for any bulk transcriptomic dataset using the tool ComboBatch that we released in the previous publication, accessible in GitHub via the link https://github.com/Nikit357/ComboBatch.

## Materials and methods

### Dataset, harmonization approaches and the analysis set

The GC B-cell lymphomas dataset, the 14 prior batch-removal strategies, the three imputation schemes, the harmonization method registry and the optional post-removal step are specified in the benchmarking study previously published by our team (Nikitin et al. 2026 Sep 17): 7,174 bulk transcriptomes of germinal-center B-cell lymphomas, pooled from 88 cohorts profiled by RNA sequencing or by Affymetrix, Illumina and Agilent microarrays. We re-used that output and computed no new harmonization. The most important algorithms were ComBat with its reference (Combat-ref) and count-based (Combat-seq) variants (Johnson et al. 2007; Zhang et al. 2020), surrogate variable analysis (Leek and Storey 2007), mutual nearest neighbors (Haghverdi et al. 2018), feature-specific quantile normalization (Franks et al. 2018), cross-platform normalization by XPN (Shabalin et al. 2008) and Shambhala-2 (Borisov et al. 2019; Borisov et al. 2022); the full registry with the parameters used is given in the source benchmark (Nikitin et al. 2026 Sep 17).

A harmonization approach is defined as one combination of batch-removal strategy, imputation, harmonization method and post-removal. We defined the approaches set by the condition that fewer than 5% of samples carry no defined expression value for all genes (pct_samples_allNA < 5). This set contained 2,234 approaches over 31 methods, 14 strategies, three imputations and both post-removal settings. Two methods, 34_arsyn and 38_harman, had a minimum of 74.23% of samples with no defined expression across all 84 of their approaches and were not included into the final approach set. Shambhala-2 was implemented using its standard P0 and Q0 calibration datasets and was named as 20_shambhala in the benchmark.

### Metric classes and the best approaches selection procedure

Of the 344 quality metrics registered by the source benchmark (Supplementary File 2 in the source article), 87 have a scoring polarity and were used for ranking in this study, and the remaining 257 were descriptive. Of the 87 scoring metrics, 51 have positive and 36 negative polarities, meaning that a higher or a lower value, respectively, is better. We assigned the scoring metrics of the source benchmark (Nikitin et al. 2026 Sep 17) to four classes by their column prefix: global, covering the 11 metrics of principal component regression (PCReg) and the dispersion separability criterion (DSC); local, covering the KNN batch-effect test (kBET), the integration and cell-type local inverse Simpson’s indices (iLISI and cLISI), average silhouette width, the cell-specific mixing score and graph connectivity; distributional, covering the Kolmogorov-Smirnov (KS) statistics and the per-gene coefficient of variation of batch means; and structural, covering tSNE and UMAP embedding entropy and centroid dispersion, the distance ratio and the Watermelon score (Supplementary File 1, sheet Metric_class_assignment). The per-component R² and variance-explained columns have no polarity, because they are descriptive and enter no class, and the seven scoring metrics that describe missing values and bimodality are likewise left unassigned. A fifth class, the composite, is the equally weighted mean of those four classes.

Within a class we converted every metric to a rank percentile over the 2,234 approaches, oriented so that a higher percentile is better according to the polarity assignments from the benchmark (Nikitin et al. 2026 Sep 17), and averaged the percentiles into one class score per approach (Supplementary File 1, sheet Class_percentile_scores). For each class we took top 5% approaches as the best ones (112 approaches in total). For comparison, we treated the 15 approaches selected by the published clustermap (Nikitin et al. 2026 Sep 17) as one more set. Overlap between each two elected sets was calculated as Jaccard index.

We also built a separate generalizability and biology preservation index as the mean of six rank percentiles: the full and multiclass-only LOBO three-class macro F1, the two-class LOBO macro F1, the mean within-cohort marker correlation before and after harmonization, the class M same-minus-different-biology margin of biomarker genes subtracted by its unharmonized baseline, and the worst multiclass fold. The worst multiclass fold (f1_mc_min) was defined as a minimum three-classes F1 score over the held-out batches with at least two biological classes. We compared batch-removal strategies against each other, imputations against each other and harmonization methods against each other, and the clustermap-selected group from the benchmark (Nikitin et al. 2026 Sep 17) was treated as an isolated named group.

### The biology evaluating metric classes L, M and N

Since gene signatures were systematically shown to predict biological subtypes better than individual genes (Jovčevska et al. 2019; Sorokin et al. 2020), for classes L and M, we took 633-gene marker panel assembled from published FL and DLBCL studies and its narrow 56-gene versions as described in the benchmarking study (Nikitin et al. 2026 Sep 17); the coverage (fraction) of each marker gene across the harmonization approaches is shown in Supplementary Figure 1A. In the L class, we calculated the Spearman correlation of these gene’s expression before and after harmonization, together with the contrast against a housekeeping control (Risso et al. 2014), and the distribution of the per-gene correlation within each signature is shown in Supplementary Figure 1C. We treated a cohort wise mean marker correlation before and after harmonization above 0.75 as a necessary condition for good quality harmonization approach.

For class M metric, we calculated Spearman rank correlation of the marker panel between pairs of batches, computed separately for pairs of the same biological group and for pairs of different groups (Nikitin et al. 2026 Sep 17). The margin was calculated as the same-biology agreement subtracted by the different-biology agreement. We also analyzed the margin relative to the unharmonized baseline expression matrices at the same strategy and imputation, so we were able to extract approaches that increased same-biology agreement and decreased different-biology agreement against their own baseline.

Class N metrics were designed as leave-one-batch-out prediction to measure a transferability and generalizability of the best harmonization approaches selected. For each approach and each of the 24 RNA batches we excluded this batch and fitted logistic regression in principal components on the training batches alone for the three-class target (DLBCL, FL and normal B cells) and for the two-class target (FL against DLBCL). Then we projected the PCA coordinates at the held-out batch and predicted its biology labels using the previously fitted logistic regression, calculating Area Under Curve (AUC) and F1 scores. We also controlled this calculation using 100 biological label permutations and averaged all the N class metrics separately on multiclass batches (having more than one biology class) and on all batches. Multiclass-only batches were selected in order to remove a potential F1/AUC bias in batches that contain single biology group. Since AUC is undefined on a held-out batch with a single biology group, AUC mean was calculated over multiclass batches only.

### Statistics and software

Group comparisons across three or more groups were done using the Kruskal-Wallis H test (Kruskal and Wallis 1952), with ε² as its effect size, and monotone associations were evaluated using Spearman correlation coefficient (Spearman 1904). Two groups comparison were performed with the two-sided Mann-Whitney U test (Mann and Whitney 1947), and matched pairs of approaches, which share strategy, method and the remaining pipeline settings and differ in one factor only, were compared with the two-sided Wilcoxon signed-rank test (Wilcoxon 1945).

Proportions of categorial variables were compared between two groups with the two-sided Fisher’s exact test (Fisher 1935), whereas proportions against the background distribution in the analysis set were evaluated using the chi-square goodness-of-fit test (F.R.S. 1900). The number of strategies in which one harshness tier ranked first was compared with the background level by the two-sided binomial test (CLOPPER and PEARSON 1934). The overlap of two elected sets was compared with the overlap expected for two random sets of the same sizes by the hypergeometric distribution (Fisher 1935), reported as the probability of an overlap at least as large, or at least as small, as observed. The p-values for comparisons were analyzed as raw, without correction for multiple testing, because individual hypothesis were tested. Per-strategy coefficients were analyzed as their ranges and medians and selected groups were reported by their medians and median absolute deviations (MAD).

We ran the analyses under Python 3.11 (3.11.16 Documentation) with pandas 2.3.3 (McKinney 2010), numpy 1.26.4 (Harris et al. 2020), scipy 1.17.1 (Virtanen et al. 2020), scikit-learn 1.8.0 (PedregosaFabian et al. 2011 Nov 1), statsmodels 0.14.6 (Seabold and Perktold 2010), matplotlib 3.10.8 (Hunter 2007) and seaborn 0.13.2 (Waskom 2021).

### AI usage

Claude Code with the Claude Opus 5 and Opus 5.5 models (The Claude 3 Model Family: Opus, Sonnet, Haiku) were used for literature retrieval, analysis code and language checks, in a plan-then-act mode: the agent wrote an implementation plan, the authors reviewed and corrected it over several cycles, and only then were the code and documents written. The study design, the choice of comparisons and the interpretation are the authors’ own.

## Results and discussion

### Metric classes capture non-overlapping components of harmonization quality

#### Global PCA-based metrics

We started the analysis from the two metric families that dominate the published comparisons, the global PCA-based and the local neighborhood and investigated their order in the canonical set of 2,234 approaches. As the global representative we took PCReg in polarity-adjusted form: 1 means that the annotation column explains none of the PCA variation and 0 means that it explains all of it. Plotting PCReg for the biology column against PCReg for the batch column gave a weak negative association across the whole set (Spearman ρ = −0.159, p = 3.7 × 10^-14^, Figure 1A and 1B for harmonization methods and batch removal strategies, respectively). The clustermap-selected approaches had a higher batch PCReg than the remaining approaches, with a median of 0.871 against 0.626 (Mann-Whitney p = 6.3 × 10^-3^). Their biology PCReg did not differ from that of the remaining approaches, with a median of 0.820 against 0.802 (p = 0.69), so the clustermap group differed from the other approaches along the batch axis of the plot only. The MADs of both comparisons are given in Supplementary File 1, sheet Paired_tests.

**Figure 1.**
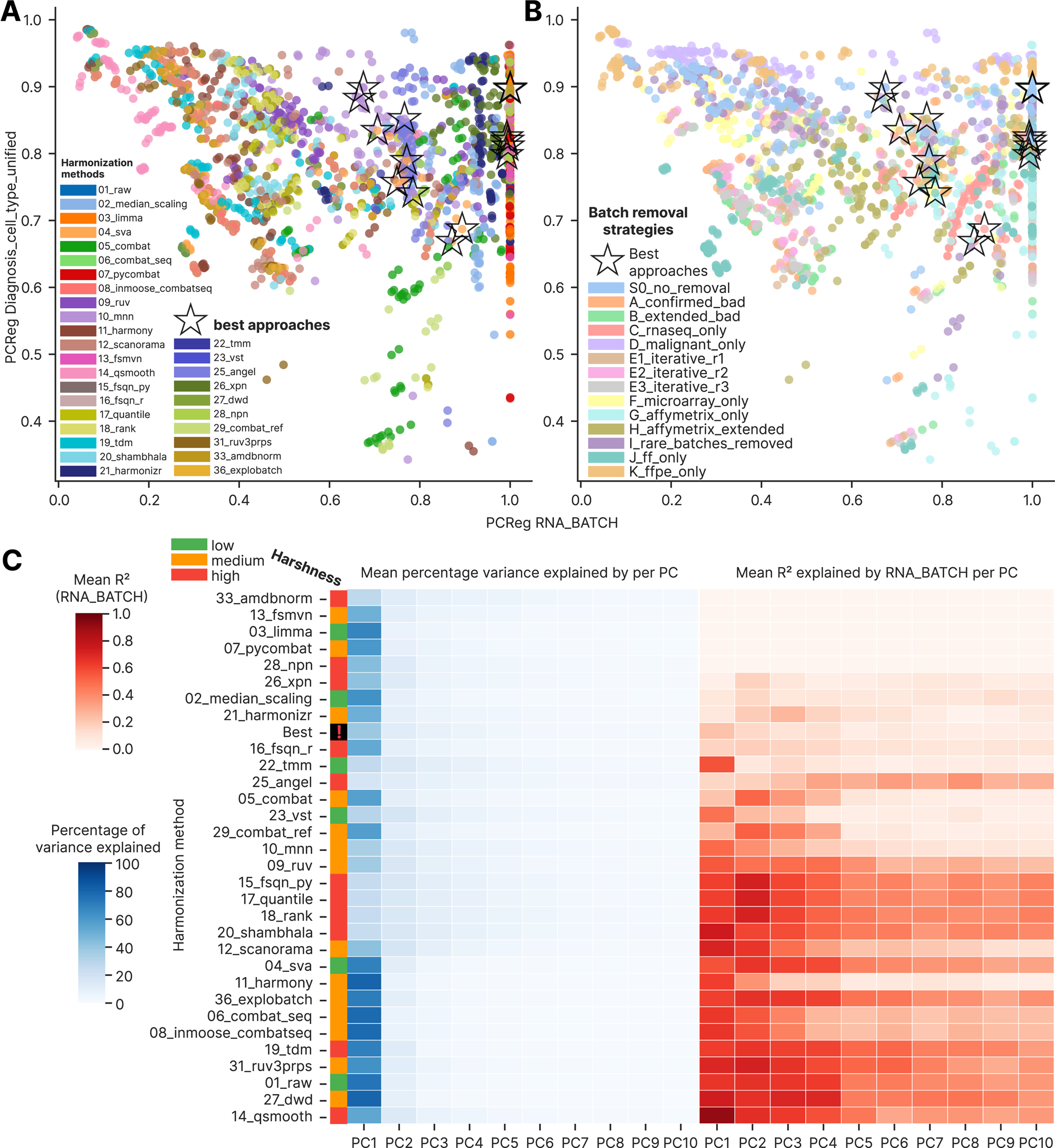
Global variance-based metrics across harmonization methods and batch-removal strategies. (A) Biology PCReg against batch PCReg for all 2,234 approaches, colored by harmonization method, with the 15 clustermap-selected approaches marked by black asterisks. (B) The same plane colored by batch-removal strategy. (C) Percentage of total variance explained by each of the first ten principal components (left) and the share of it explained by the batch column (right), grouped by method and ordered by mean batch PCReg. The clustermap group is shown as one composite row.

Individual methods behaved in divergent and counterintuitive manner. The batch PCReg of 33_amdbnorm was exactly 1.000 in each of its 14 completed runs. The method 13_fsmvn reached a median of 1.000 with minimum 0.997, but both methods were selected as the clustermap best approaches only for the no-removal strategy (Figure 1A). The methods 05_combat and 29_combat_ref separated biology better than the remaining methods across the A, B, I, J and E1 to E3 strategies (median biology PCReg 0.568 against 0.778, Mann-Whitney p = 1.3 × 10^-27^, Figure 1A) without being selected by the clustermap. Their biology PCReg spanned 0.357 to 0.823 over these strategies. By PCA batch correction, 14_qsmooth was the last method, with a median of 0.208 against 0.672 for the other methods (p = 1.3 × 10^-28^; its MAD is given in Supplementary File 1, sheet Paired_tests). Among strategies, malignant-only, FFPE-only and no-removal were weakest on biology, with medians of 0.939, 0.925 and 0.883, respectively. Each of the three was higher than the other strategies (Mann-Whitney p ≤ 6.4 × 10^-27^). RNA-seq-only was strongest on batch PCReg, with a median of 0.872 against 0.563 for the other strategies (p = 4.5 × 10^-20^, Figure 1B). The malignant-only and FFPE-only strategies contain no normal B cells, so their biology column had only four classes (DLBCL, FL, Burkitt lymphoma and High-grade B-cell lymphoma) potentially explaining their high PCReg by biology.

Decomposing the variance by principal component showed the batch signal transformation by harmonization methods (Figure 1C; Supplementary Figure 2A, B). Five methods (15_fsqn_py, 17_quantile, 18_rank, 20_shambhala and 25_angel) spread the variance slowly across components, with 17.3 to 25.3% on the first component. On the second component the same five methods placed 11.3 to 15.3%. Eight further methods and the clustermap best group concentrated 26.2 to 43.3% on the first component (Figure 1C). The remaining 18 methods had up to 79.9% of variance in the first component, with the maximum for 11_harmony and 79.5% for 27_dwd. For the five leading methods by mean batch PCReg (03_limma, 07_pycombat, 13_fsmvn, 28_npn and 33_amdbnorm) the batch-explained R^2^ stayed below 1% at every one of the first ten components, with a maximum of 0.002 at the second component of 07_pycombat (Supplementary Figure 2B). Without harmonization and without batch removal the same dataset gives a batch PCReg of 0.226, which means that batch accounts for 77.4% of the PCA variance summed over all components.

#### Local metrics

The local metrics ordered the approaches in a different manner compared to the global metrics (Figure 2). Taking the local inverse Simpson’s index as a main local metric, 10_mnn had the maximum batch mixing with a median iLISI of 1.72 against 1.16 for the remaining approaches (Mann-Whitney p = 3.3 × 10^-40^). 14_qsmooth showed the lowest mixing, at a median of 1.07 (p = 8.0 × 10^-27^ against the rest, Figure 2A). Batch and biology mixing increased together in a trade-off pattern (Spearman ρ = 0.527, p = 7.0 × 10^-160^, Figure 2A). If stratified by strategy, the association was confirmed inside every strategy except the Affymetrix-only one, with ρ between 0.374 and 0.815 (each p ≤ 9.2 × 10^-7^, Figure 2B; Supplementary File 1, sheet Scatter_correlations). Inside the Affymetrix-only strategy it was not significant (ρ = 0.147, p = 0.069). The tSNE entropy of the batch labels agreed with the methods LISI ordering, with a median of 0.179 for 10_mnn against 0.018 for 14_qsmooth (Mann-Whitney p = 2.3 × 10^-28^, Figure 2C; the MADs are in Supplementary File 1, sheet Paired_tests). In contrast, tSNE centroid dispersion, a global quantity computed in the same tSNE embedding, split the methods into a low group and a high group, with method means between 0.24 for 13_fsmvn and 1.00 for 27_dwd, and had the clustermap group on the boundary between them (Figure 2D).

**Figure 2.**
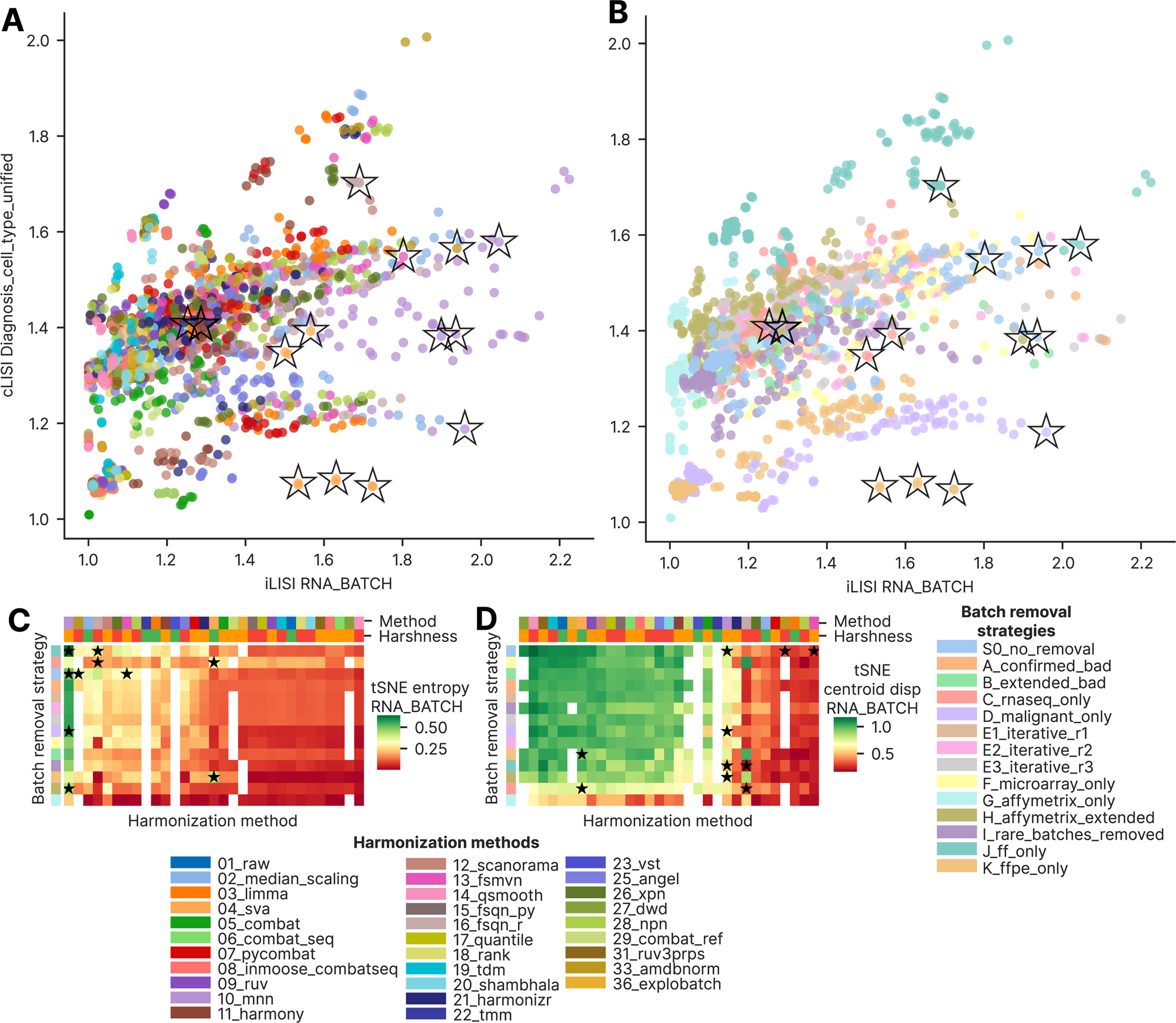
Local neighborhood metrics across harmonization methods and batch-removal strategies. (A) Biology cLISI against batch iLISI for all 2,234 approaches, colored by method. (B) The same plane colored by strategy. (C) Heat map of tSNE entropy of the batch labels by method and strategy. (D) Heat map of tSNE centroid dispersion of the batch labels over the same grid. Clustermap-selected approaches are marked by black asterisks throughout.

#### Distributional metrics

The third family, distributional convergence, separated the approaches on a different axis compared to global and local metrics. We compared the mean within-batch KS D statistic over 1,000 randomly drawn genes against the mean between-batch D statistic for the same genes, and the two metrics were unrelated (Spearman ρ = 0.051, p = 0.021). Individual methods formed vertical bands, and strategies formed horizontal layers (Figure 3A, 3B, respectively). The clustermap group had a lower within-batch D than the remaining approaches, with a median of 0.445 against 0.577 (Mann-Whitney p = 1.3 × 10^-4^). It’s between-batch D was lower as well, with a median of 0.386 against 0.597 (p = 2.3 × 10^-3^). The RNA-seq-only strategy located at the lower left of the plane, with a within-batch median of 0.439 against 0.581 for the other strategies (p = 2.1 × 10^-79^, Figure 3B). It’s between-batch median was 0.332 against 0.628 (p = 2.0 × 10^-43^). The MADs of these comparisons are again given in Supplementary File 1, sheet Paired_tests.

**Figure 3.**
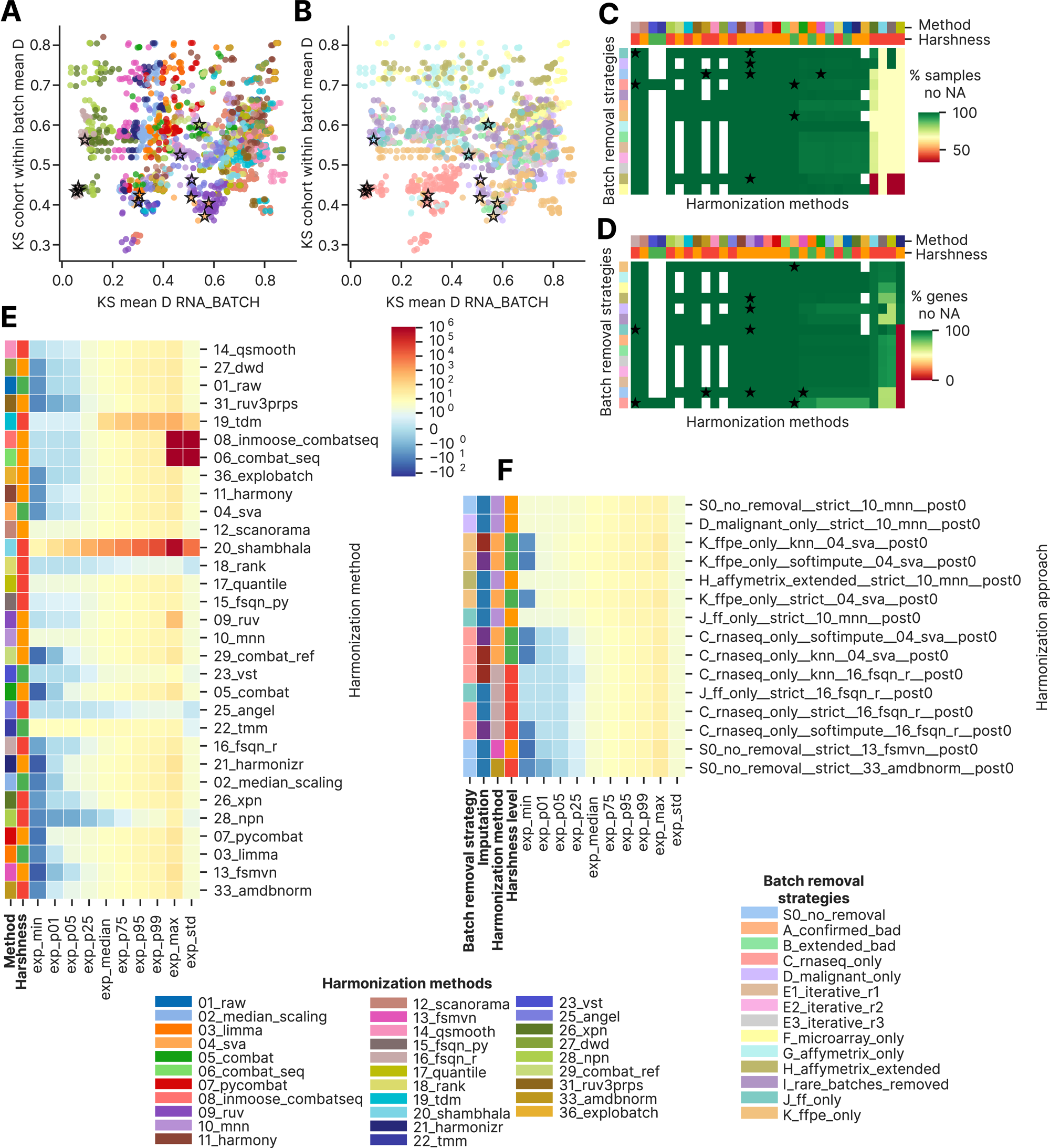
Distributional convergence, missing values and output scale. (A) Mean between-batch against mean within-batch Kolmogorov-Smirnov D statistic over 1,000 randomly drawn genes, colored by method. (B) The same plane colored by strategy. (C) Heat map of the percentage of samples with no missing expression value, by method and strategy. (D) The same heat map for genes. Clustermap-selected approaches are marked by black asterisks in panels A to D. (E) Heat map of the minimum, the 1st, 5th, 25th, 50th, 95th and 99th percentiles, the maximum and the per-gene standard deviation of the output, averaged by method and ordered by mean batch PCReg. (F) The same heat map for the 15 clustermap-selected approaches.

#### Gene expression scale and missingness metrics

Missing values and normalized gene expression scale separated the methods differently again. Of the 31 methods, 18 lost no sample to a missing value under any strategy (Figure 3C). Nine methods lost 0.003 to 1.4% of samples due to all their gene expressions being non-defined. Four methods (26_xpn, 20_shambhala, 15_fsqn_py and 17_quantile) lost 30.2 to 38.1% samples. At gene level the same 18 methods lost no genes (Figure 3D). Ten methods lost 0.004 to 2.41% of genes with their expression being non-defined in all samples. Furthermore, 15_fsqn_py lost 13.0% and 17_quantile lost 16.2%. The method 21_harmonizr lost all genes under eight of the 14 strategies.

At the level of harmonized gene expression scale, the methods 06_combat_seq and 08_inmoose_combatseq returned maximum expression values of 2.76 × 10^8^ and 1.46 × 10^11^. Their 99th percentiles were 18.5 and 17.9, indicating 6 to 9 orders of magnitude derived from outliers. Averaged over its 84 approaches, 20_shambhala returned a median expression value of 1,359.6 and a maximum of 555,004.7. Of the 31 methods, 19 returned a minimum below zero, with 13_fsmvn lowest at −40.6 (Figure 3E). Across the clustermap best approaches the median expression varied from 2.63 to 6.70 (Figure 3F). Their maximum expression varied from 12.6 to 28.6.

Taken together, the initial screening of harmonization approaches by global, local and distributional metrics showed that the three families ordered the same set of approaches differently, indicating potential trade-offs between these metric classes. In the next chapters we deeply investigated these differences in order to derive a unified framework of harmonization approaches benchmarking.

### Global variance-based metrics saturate below the threshold of biological preservation

#### Global PCA-based metrics in-depth comparison

Twelve of the 31 methods overlapped batches better than biology in PCA space, and five of them (03_limma, 07_pycombat, 13_fsmvn, 28_npn and 33_amdbnorm) reached a mean batch PCReg at least 0.999, with 33_amdbnorm at exactly 1.000 (Supplementary Figure 3A). At strategy level only two of the 14 strategies, RNA-seq-only and Affymetrix-only, overlapped batches better than biology (Supplementary Figure 3B). Batch PCReg differed across methods far more than across strategies (Kruskal-Wallis ε² = 0.739 against 0.100, both p ≤ 6.7 × 10^-43^), which indicates superiority of the choice of method at this level of batch effect quality control.

Looking at the PCReg-saturated approaches, we examined the 198 approaches whose batch PCReg exceeded 0.9999, which were generated by six methods (03_limma, 13_fsmvn, 07_pycombat, 21_harmonizr, 33_amdbnorm and 28_npn). Of these, 98.5% returned a kBET acceptance rate under 0.1 (Supplementary Figure 4E, F). Over all approaches the same share was 97.0%, and the saturated approaches did not differ from the remaining ones (96.9%; Fisher’s exact p = 0.27). PCReg saturation therefore does not indicate poorer local batch mixing than the majority of the benchmark.

#### Global PCA-based metrics by other columns and Dispersion Separability Criterion

The remaining global metrics closely reproduced PCReg and added a minor separation. We first compared PCReg for the batch column with PCReg for the platform and cohort columns (Supplementary Figure 5A, B, respectively). Because one platform can correspond to two or three batches, PCReg by platform followed PCReg by batch almost linearly (Spearman ρ = 0.974, p < 10^-300^, Supplementary Figure 5A), with the FFPE-only and FF-only strategies closest to the diagonal, where each platform keeps a single batch. PCReg by cohort followed PCReg by batch less closely (ρ = 0.606, p = 2.1 × 10^-224^), and most approaches retained more cohort-related than batch-related variance (Supplementary Figure 5B).

DSC for the batch column followed PCReg along a monotone sigmoid (ρ = −0.970, p < 10^-300^, Supplementary Figure 5C). DSC for the platform column followed DSC for the batch column linearly on a logarithmic scale (ρ = 0.966, p < 10^-300^, Supplementary Figure 5D). DSC for the cohort column, in contrast, formed two clouds of points that the strategies did not explain and that more plausibly reflect the harmonization method (ρ = 0.544 against batch DSC, p = 1.6 × 10^-172^, Supplementary Figure 5E). All 15 clustermap approaches were below the overall median batch DSC of 0.962 and lower than the remaining approaches (Mann-Whitney p = 0.011). The four MNN approaches of the group were located at the boundary (0.781 to 0.957). The FSQN R approaches had the metric about two orders of magnitude lower (0.001 to 0.010). Such behavior can be explained by the definition of DSC as between-group scatter divided by within-group scatter, which is sensitive to outliers.

#### tSNE Centroid Dispersion and Average Silhouette Width

We also studied tSNE centroid dispersion by harmonization approaches. By batch this metric decreased with batch PCReg along an arc (ρ = −0.864, p < 10^-300^, Supplementary Figure 5F, G by strategies and methods, respectively). A subgroup of 11_harmony approaches under the FF-only and FFPE-only strategies were situated below this trend, although their dispersion did not differ from that of the other methods within either strategy (Mann–Whitney p = 0.32 and 0.15, respectively). Linking each approach with and without post-removal showed that post-removal raised batch PCReg under every one of the 14 strategies (Wilcoxon signed-rank, p ≤ 4.3 × 10^-6^ for each; Supplementary Figure 5H). The shift was largest in the Affymetrix-only strategy, with a median of 0.075 against 0.007 to 0.028 elsewhere, presumably because post-removal drops the already quantile-normalized SOM cohort, which makes up a large share of this strategy.

Average silhouette width tracked PCReg in a noisier manner (ρ = −0.630, p = 1.9 × 10^-247^, Supplementary Figure 5I, J by strategies and methods, respectively). The Affymetrix-only and FFPE-only strategies were worst by the metric, with means of 0.165 and 0.077, each above the other strategies (Mann-Whitney p ≤ 3.8 × 10^-7^, Supplementary Figure 5I). The best methods were 02_median_scaling and 33_amdbnorm, with means of −0.286 and −0.276 (p ≤ 1.3 × 10^-8^ against the rest, Supplementary Figure 5J).

#### Impact of imputation and batch removal strategy on PCReg

The two pipeline factors, namely imputation and post-removal, impacted the harmonization performance to the least extent according to the source benchmark (Nikitin et al. 2026 Sep 17), and we quantified them here on the global metric. We compared the imputations over the 11 methods with the lowest batch PCReg (Supplementary Figure 6A). Softimpute exceeded KNN imputation in these methods by between 1.7 and 9.7 percentage points, with a mean of 4.8. In matched pairs of approaches, which share strategy, method and post-removal, the median gain of softimpute over KNN imputation was 0.042 in batch PCReg (Wilcoxon signed-rank p = 1.5 × 10^-28^). Strict exclusion scored below KNN imputation in the same matched comparison, by a median of 0.020 (p = 2.7 × 10^-22^). Over the 11 methods, the means were 0.375 for strict exclusion against 0.394 for KNN imputation. Post-removal raised the mean batch PCReg for 30 of the 31 methods (Supplementary Figure 5H). Over matched pairs of approaches the median shift was 0.011 (Wilcoxon signed-rank p = 1.0 × 10^-140^). The largest increase was for 22_tmm, from 0.70 up to 0.88. The next method was 23_vst, which increased from 0.70 to 0.83, both methods were defined for RNA-seq only strategy. The single method with PCReg decrease was 12_scanorama, from 0.49 to 0.48 (Supplementary Figure 6C).

As it was shown previously (Nikitin et al. 2026 Sep 17), batch removal strategies impacted batch PCReg to higher degree compared to imputations and post removals (Kruskal–Wallis ε² = 0.100 across strategies against 0.002 across imputations and 0.002 across post-removal settings). Every method reached at least 0.76 under the RNA-seq-only strategy, 0.72 under Affymetrix-only and 0.46 under Affymetrix-extended (Supplementary Figure 6B). In contrast, the 16_fsqn_r method decreased from its cross-strategy mean of 0.83 to 0.30 under microarray-only and 0.08 under FFPE-only. Under the S0 no removal strategy the same method reached a median of 0.995, and both strategies were below it (Mann-Whitney p = 2.2 × 10^-3^ for each), proving that feature-specific quantile normalization performs poorly in cross-platform FFPE-only dataset.

#### Principal components in-depth comparison by strategies of batch removal

As defined above, a PCReg of 0 is the worst scenario for the batch columns (RNA_BATCH, PLATFORM_RNA, COHORT_LABEL, RNASEQ_SOURCE) and the best one for the biology columns (Major_Group, Diagnosis_cell_type_unified, TUMOR_NORMAL). Because the variance is unevenly distributed across the PCs, we compared PCReg RNA_BATCH against R^2^ of RNA_BATCH in the first PC by harmonization methods and batch removal strategies (Supplementary Figure 6D, 6E, respectively). As expected, we observed a strong negative relationship between the two (Spearman ρ = −0.939, p < 10^-300^), with the clustermap best approaches occupying the lower right corner of the plot. 10_mnn, 20_shambhala and 16_fsqn_r were shifted above the main diagonal, suggesting that their batch-explained variance is moved towards lower rank PCs. The same was true for the RNA-seq-only strategy (Supplementary Figure 6E).

For all the batch removal strategies we also investigated batch-explained variance for the first 10 PCs (Supplementary Figure 6F; Supplementary Figure 2C, D), examining the percentage of variance explained and the R^2^ for RNA_BATCH at each PC. The joint group of the clustermap best approaches ranked second by batch PCReg after the RNA-seq-only strategy (shown earlier in Supplementary Figure 3B). Both the RNA-seq-only strategy and the clustermap best approaches group showed the lowest average percentage of variance explained by the first PC (30.4 and 37.9%, respectively), and the smoothest decay of the percentage by PC number (Supplementary Figure 2C). The RNA-seq-only value was lower than over the other strategies (Mann-Whitney p = 4.7 × 10^-36^). The clustermap group value was lower than over the remaining approaches (Mann-Whitney p = 0.038). The Affymetrix-only strategy showed the highest average percentage, 66.0% (Mann-Whitney p = 6.1 × 10^-16^ against the other strategies), presumably because the already quantile-normalized SOM cohort makes up a larger share of the samples in this strategy than in any other. The decay trajectory of the RNA_BATCH R^2^ for the RNA-seq-only strategy and the best approaches started at 0.29 and 0.23, respectively, whereas the other strategies, except the Affymetrix-only and Affymetrix-extended ones, had higher first-PC values (Supplementary Figure 2D). Of the 14 batch removal strategies, 10 had a lower RNA_BATCH R² for the first PC than for the second.

Taking together, global metrics in-depth analysis showed that global metrics form a weak, permissive criterion of harmonization quality. Both biology and batch PCReg should be measured, and other metric classes should be added to evaluate harmonization quality robustly. Moreover, different groups of metrics (PCReg, tSNE centroid dispersion, silhouette width) select different best approaches. Therefore, event within global metrics there is no agreement about the best harmonization approaches which emphasizes the necessity of other metric classes in a benchmarking framework.

### Batch-associated variance is redistributed across principal components instead of being eliminated

We separately investigated distribution of batch-associated variance over principal components by harmonization methods and batch removal strategies (Supplementary Figure 2A–D). Ten of the 14 strategies, and 19 of the 31 methods, had a lower batch-explained R^2^ on the first principal component than on the later components (Supplementary Figure 2B, D). Among the five methods whose explained variance decayed most slowly, four (excluding 20_shambhala) reached their maximum batch R^2^ away from the first component (Supplementary Figure 2B). On the first component, batch explained 22.9% of the variance in the clustermap group, 64.8% in the unharmonized data and 90.8% under 14_qsmooth, the weakest method on the PCA metrics (Supplementary Figure 2B). The first component itself explained 37.9%, 73.5% and 53.7% of the total variance in these three groups, respectively (Supplementary Figure 2A). On the second component the batch R^2^ was 15.2%, 65.9% and 67.1%, respectively, so the unharmonized data and 14_qsmooth kept most of their batch signal beyond the first component. The second component itself explained 11.6%, 9.7% and 13.5% of the total variance in the clustermap, unharmonized and 14_qsmooth.

The individual principal components analysis therefore showed that harmonization benchmarking based on the first component only inherently underestimates batch effect, and harmonization methods can hide the batch-related variance into the lower rank principal components which should be accounted for during benchmarking.

### Local neighborhood metrics discriminate well between good and bad approaches but are vulnerable to batch composition

#### The main local metrics: LISI, kBET, graph connectivity

We extensively compared local neighborhood metrics between harmonization methods and batch removal strategies. Only six of the 31 methods (02_median_scaling, 10_mnn, 16_fsqn_r, 25_angel, 28_npn and 33_amdbnorm) mixed batches better than biology by LISI (Supplementary Figure 7A). The kBET acceptance rate behaved as more stringent criterion: it remained under 0.01 for 82.0% of the 2,234 approaches and under 0.10 for 97.0% of them (Supplementary Figure 8A). Five clustermap best approaches exceeded 0.10, against 2.8% of the remaining approaches (Fisher’s exact p = 5.0 × 10^-5^), and all five were SVA approaches under the RNA-seq-only and FFPE-only strategies. Eight methods exceeded the batch kBET acceptance rate of the unharmonized baseline (0.027) and only two of them did so with a relatively wide margin, 22_tmm (0.331) and 23_vst (0.230), both run under the RNA-seq-only strategy alone (Supplementary Figure 8C).

According to their design, local values depended on how many classes the strategy leaves in the data, which limits comparison across strategies (Korsunsky et al. 2019b). It was therefore expected that the malignant-only and FFPE-only strategies retained four malignant groups and returned the lowest biology LISI (medians 1.10 and 1.07). The FF-only strategy preserved all the normal B-cell types and returned the highest value (median 1.62). Each of the three strategies differed from the other strategies significantly (Mann-Whitney p ≤ 2.8 × 10^-76^, Supplementary Figure 7B). The same two strategies returned the lowest biology graph connectivity, with means of 0.727 for FFPE-only and 0.646 for malignant-only (each lower than the other strategies, Mann-Whitney p ≤ 6.0 × 10^-24^, Supplementary Figure 9D), although fewer biology groups are expected to increase connectivity, which could derive from their assembly from heterogeneous FFPE batches.

Graph connectivity by biology and by platform were nearly unrelated (Spearman ρ = 0.080, p = 4.3 × 10^-4^, colored by method in Supplementary Figure 9C). In this plot 10_mnn, 25_angel, 09_ruv and 16_fsqn_r were located in the upper right corner near the clustermap best approaches, and 02_median_scaling, 07_pycombat, 03_limma, 04_sva and 11_harmony in the lower left corner, with both lower biology preservation and a more fragmented platform-level structure (Supplementary Figure 9C). The RNA-seq-only strategy produced a tight group of 41 approaches with kBET acceptance rates between 0.654 and 0.867 (Supplementary Figure 8B). These approaches originated from 18 methods, and none of them was selected by the clustermap, because their LISI values remained between 1.24 and 1.38.

We then visualized batch and biology LISI on heatmaps by harmonization method and batch removal strategy (Supplementary Figure 7C, D, respectively). The clustermap best approaches occupied local peaks of batch LISI (Supplementary Figure 7C) but not the corresponding local minima of biology LISI (Supplementary Figure 7D). Against the other approaches, the clustermap group had a higher batch LISI (Mann–Whitney p = 2.0 × 10^-7^). Its biology LISI was indistinguishable from theirs (p = 0.80), which reflects how small the biological differences are relative to the technical ones in this dataset both at the global and local level. By strategy, the RNA-seq-only strategy had the highest batch kBET acceptance rate, followed by the two Affymetrix strategies and the FF-only strategy, whereas removing bad or rare batches, or keeping malignant samples only, lead to kBET score near zero (Supplementary Figure 8D; Kruskal–Wallis across strategies p = 1.2 × 10^-78^).

#### tSNE and UMAP based local metrics

We also compared LISI with the embedding-based mixing metrics. The tSNE entropy of the batch labels followed batch LISI closely (Spearman ρ = 0.915, p < 10^-300^), and 10_mnn together with the three clustermap 04_sva approaches formed a separate linear trend with a higher entropy for a given LISI (Supplementary Figure 8E). By strategy the differences were smaller, and the S0 no-removal, rare-batches-removed, Affymetrix-extended and microarray-only strategies were located on the lower side of the trend (Supplementary Figure 8F).

tSNE centroid dispersion decreased as tSNE entropy rose (ρ = −0.493, p = 5.6 × 10^-137^, Supplementary Figure 8G, H). 10_mnn, with a minority of the 04_sva, 11_harmony, 12_scanorama and 16_fsqn_r approaches, formed a diagonal cloud of high entropy, whereas the other methods formed two dense, round clouds (Supplementary Figure 8G). The strategies formed layers present in both clouds (Supplementary Figure 8H), so a given method can produce either good or poor batch overlap in the embedding, and the composition of the batches governs how well they mix locally. UMAP and tSNE entropy agreed almost exactly (ρ = 0.992, p < 10^-300^) and added no difference between methods or strategies (Supplementary Figure 8I, J, respectively). UMAP and tSNE centroid dispersion were also strongly correlated (ρ = 0.814, p < 10^-300^), with the same two clouds of methods (Supplementary Figure 8K) and each strategy present in both (Supplementary Figure 8L). The clustermap best approaches were spread along this trend line, so the selection gave priority to local mixing over group separation in both non-linear embeddings.

#### Local and global PCA metrics

Finally, we compared LISI and kBET against PCReg (Supplementary Figure 4). Batch LISI increased with batch PCReg (Spearman ρ = 0.557, p = 1.4 × 10^-182^), with 10_mnn, 04_sva, 11_harmony, 12_scanorama and 16_fsqn_r above the general trend (Supplementary Figure 4A). By strategy the approaches formed layers similar to, but noisier than, those of tSNE centroid dispersion against tSNE entropy (Supplementary Figure 4B). For the biology column the relationship was negative (ρ = −0.403, p = 4.0 × 10^-88^), and the clustermap best approaches were spread uniformly over the plane (Supplementary Figure 4C). Strategies formed horizontal layers of variable PCReg and fixed biology LISI (Supplementary Figure 4D), the malignant-only and FFPE-only layers at the lowest LISI and the FF-only layer at the highest, in line with the number of biological classes each strategy keeps. kBET acceptance rate increased weakly with batch PCReg (ρ = 0.396, p = 1.1 × 10^-84^, Supplementary Figure 4E), and only 04_sva and 10_mnn had a kBET above 0.1 in a sizable number of their approaches. By strategy, the RNA-seq-only approaches formed a compact group of high kBET and high PCReg (Supplementary Figure 4F), none of which was selected by the clustermap.

Taking together, the in-depth analysis of local metrics showed that reaching the maximum of one local metric is not sufficient for the clustermap best selection. Moreover, the local metrics were highly dependent on prior batch removal strategies. These findings indicate that a harmonization quality benchmarking pipeline should treat local metrics as necessary but not sufficient metric family.

### Distributional convergence is the only axis on which the high harshness harmonization methods rank first

#### Kolmogorov-Smirnov test based metrics

We further deeply compared metrics of distributional similarity between harmonization approaches. On the fraction of cohort pairs whose expression distributions differ by the KS test, 29 of the 31 methods had a mean score below the unharmonized baseline of 0.948 (Supplementary Figure 9A). By metric ranks, 26 methods had lower values than the baseline (two-sided Mann-Whitney p < 0.05 each), and 04_sva, 31_ruv3prps and 36_explobatch did not differ from it (Supplementary File 1, sheet Paired_tests). Only 27_dwd performed poorer than the raw expression matrix, at 0.965 but without significant differences (two-sided Mann-Whitney p = 0.46). Only ten methods showed the mean below 0.7, ranging from 0.173 for 28_npn to 0.691 for 03_limma. The clustermap best group reached a mean of 0.560, lower than the remaining approaches (Mann-Whitney p = 9.6 × 10^-3^), so 56.0% of cohort pairs still differed after the best harmonization by the clustermap assessment. For the strategies comparison, the S0 no-removal strategy had the lowest KS fraction significant, followed by the clustermap group and the RNA-seq-only strategy (no removal against the other strategies, Mann–Whitney p = 0.0033; Supplementary Figure 9B).

#### Within to between batch distance ratio

For another metric, ratio of within-batch to between-batch distances, we observed separation of the clustermap group as well, with a median batch distance ratio of 0.914 against 0.757 for the remaining approaches (Mann-Whitney p = 1.0 × 10^-3^, Supplementary Figure 9E). Only the two methods 27_dwd (0.508) and 14_qsmooth (0.531) had the metric below the unharmonized dataset (0.566). The batch and biology distance ratios were nearly unrelated (Spearman ρ = −0.074, p = 4.8 × 10^-4^). By strategy, the malignant-only, no-removal and FFPE-only strategies had the highest biology distance ratio and the weakest biology preservation (Kruskal–Wallis across strategies p = 5.8 × 10⁻¹⁶¹; Supplementary Figure 9F).

#### Watermelon score

We then analyzed the Watermelon score, which measures how well an approach separates classes on a clustermap in gene expression space, by batch and by biology columns (Supplementary Figure 9G–J). On the batch column the clustermap group had a lower Watermelon score than the remaining approaches, a median of 0.318 against 0.352 (Mann-Whitney p = 5.0 × 10^-4^, Supplementary Figure 9G). On the biology groups there were no difference (0.290 against 0.306, p = 0.086). The batch and biology scores were moderately correlated (Spearman ρ = 0.422, p = 5.0 × 10^-97^, Supplementary Figure 9G). The lowest batch score belonged to 10_mnn and three RNA-seq-only 11_harmony approaches (Supplementary Figure 9G). By strategy, the Affymetrix-only strategy scored worst on the batch column, and the malignant-only and FFPE-only strategies, with four malignant classes each, scored best on the biology column (Supplementary Figure 9H). Averaged over all batch columns, the clustermap group again scored lower than the remaining approaches (median 0.311 against 0.368, p = 1.4 × 10^-5^, Supplementary Figure 9I). Averaged over the biology columns, the clustermap group did not differ from them (median 0.350 against 0.372, p = 0.21). The two averages, the batch and the biology, were weakly related (ρ = 0.124, p = 3.6 × 10^-9^, Supplementary Figure 9I), with 10_mnn forming a compact group of low batch scores. The strategies separated into compact groups in the two average WM coordinates, the malignant-only approaches lowest on the biology average and the FFPE-only approaches highest, although both keep the same four malignant classes (Supplementary Figure 9J). Because the Watermelon score separated the clustermap group on the batch columns only, its biology component could capture a mode of harmonization quality that the clustermap selection did not emphasize, which makes this metric a candidate for an independent metric class to be compared to the global and local families.

Additionally, we compared output expression quantiles by strategies and found that they were similar, except for the Affymetrix-only strategy, which had a larger per-gene standard deviation than the other strategies (median 3.60 against 2.75, Mann–Whitney p = 6.8 × 10^-6^; Supplementary Figure 9K). Its maximum expression, by contrast, did not differ from the other strategies (p = 0.30).

#### Comparison of methods by their harshness score

We then compared methods by their harshness score defined in the harmonization benchmark (Nikitin et al. 2026 Sep 17) as low harshness is linear transformation, high harshness is significant distribution reshaping (for example in quantile normalization) and medium harshness is everything in between. There were 344 low, 1,042 medium and 848 high harshness approaches for the comparison. Within each strategy we identified the tier with the highest or the lowest mean of a given metric, and compared the number of strategies in which the same tier ranked first with the one-in-three chance level by the two-sided binomial test.

Low-harshness methods had the highest mean batch PCReg in 11 of the 14 strategies (binomial p = 6.9 × 10^-4^, Supplementary Figure 10A). They also had the highest mean batch LISI in 10 of 14 strategies (p = 4.0 × 10^-3^, Supplementary Figure 10D). High-harshness methods had the lowest tSNE entropy in 12 of 14 (p = 8.2 × 10^-5^, Supplementary Figure 10C). High harshness prevailed over low and middle harshness on one axis only, namely distributional convergence, where it demonstrated the lowest KS fraction significant in 13 of the 14 strategies (p = 6.1 × 10^-6^, Supplementary Figure 10F). Inside any single strategy, the tier difference reached significance in at most two of the 14 strategies these metrics (Kruskal-Wallis p < 0.05), so the finding relies on the consistent direction across strategies, while the difference inside one strategy is relatively small. On the biology columns, low-harshness methods had the highest biology LISI (the weakest local preservation of biology), in 12 of the 14 strategies (binomial test against p = 8.2 × 10^-5^; Supplementary Figure 10E), so their stronger local mixing of batches is connected with weaker local preservation of biology. On biology PCReg the low tier was lowest in 7 of the 14 strategies, which does not exceed the random expectation (p = 0.25; Supplementary Figure 10B). Matching expression distributions to a same distribution is a distinguishing feature of the harshest transformations, but it does not lead to improvement of local or global metrics.

### Composite score of the 87 polarity metrics supports the clustermap best approaches

We also calculated and compared a composite score, which was the equally weighted composite of all the 87 metrics. Expectedly, it ranked the clustermap group as first at a mean of 0.602, above the remaining approaches (Mann-Whitney p = 1.8 × 10^-8^, Supplementary Figure 11A). The RNA-seq-only strategy was next, at a mean of 0.578 (p = 1.6 × 10^-55^ against the other strategies). To find which metrics drive the composite, we split the score of each approach into the contributions of each metric type, metric group and annotation column, and measured the spread of each contribution as the range of its group means (maximum minus minimum), counting the 15 clustermap-selected approaches as one additional group (Supplementary File 1, sheet Composite_dispersion). By metric type of the source benchmark, global distance metrics provided 47 of the 87 columns and 0.360 of the clustermap group’s score (Supplementary Figure 11A, B). Their contribution ranged by 0.066 across strategies and by 0.085 across methods. Local neighborhood metrics supplied only 22 columns and 0.109 of the score (to the clustermap best approaches). Their contribution nonetheless ranged by 0.071 across strategies and by 0.075 across methods, a spread close to that of the global distance metrics from less than a third of their absolute contribution. This finding indicates that less groups of global metrics (for example, PCReg only) should be taken into harmonization quality benchmarks for future evaluations.

By metric group in the composite score, the PCA-based variance decomposition metrics (PCReg and DSC) ranged by only 0.017 across strategies and 0.030 across methods (Supplementary Figure 11C, D, respectively). The neighbor-based integration metrics ranged by 0.043 across strategies and 0.049 across methods. By annotation column, batch columns supplied 57 of the 87 metrics and 0.403 of the clustermap group’s score (Supplementary Figure 11E, F). The 18 biology columns separated approaches to lesser degree than the batch columns did: their contribution ranged by 0.053 across strategies, against 0.121 for the batch columns (Supplementary Figure 11E). Across methods the two ranges were 0.041 and 0.154, respectively (Supplementary Figure 11F).

We also sorted all approaches by the composite score and revealed an upper tail of 150 approaches above 0.6, which is 6.7% of all approaches (Supplementary Figure 12). The tail contained nine of the 15 clustermap selections, against 6.4% of the remaining approaches (Fisher’s exact p = 7.8 × 10^-8^). We did not reveal ordering of the harshness tiers by this composite score (Kruskal-Wallis p = 0.48).

### Biology preservation metrics behavior by harmonization approaches: harsh methods partially corrupt biology but predict biology classes better

We then compared the biology related metrics of classes L, M and N between the harshness groups and observed significant differences (Table 1; Supplementary Figure 13A–L). Each metric was compared across the three tiers by the Kruskal-Wallis H test and between each pair of tiers by the two-sided Mann-Whitney U test, and the number of approaches behind each metric is shown in Supplementary File 1, sheet Harshness_full. Low-harshness methods kept the within-cohort marker correlation close to its maximum (Supplementary Figure 13A) and had the smallest marker-minus-housekeeping contrast (Supplementary Figure 13B), which was expected under the minimum linear or quasi-linear data transformations.

**Table 1.** Classes L, M and N by method harshness tier. Mean and median over the approaches of each tier (low, medium, high); Kruskal-Wallis H and p across the three tiers; two-sided Mann-Whitney p for each pair of tiers. All p-values are raw. LOBO3, three-class target (DLBCL, FL, normal B cells); LOBO2, two-class target (FL against DLBCL). AUC is undefined on a single-class held-out batch and is therefore reported on multiclass folds only. The number of approaches per metric and tier is given in Supplementary File 1, sheet Harshness_full.

| Class | Metric | Fold cut | Low, mean / median | Medium, mean / median | High, mean / median | Kruskal-Wallis H | p | p low-medium | p low-high | p medium-high |
| --- | --- | --- | --- | --- | --- | --- | --- | --- | --- | --- |
| L | Mean within-cohort marker correlation | n/a | 0.982 / 0.998 | 0.746 / 0.880 | 0.927 / 0.922 | 340.1 | $1.4 \times 10^{-74}$ | $3.7 \times 10^{-51}$ | $9.2 \times 10^{-84}$ | $5.8 \times 10^{-5}$ |
| L | Marker minus housekeeping correlation | n/a | 0.010 / -0.000 | 0.052 / 0.009 | 0.029 / 0.031 | 139.3 | $5.5 \times 10^{-31}$ | $5.5 \times 10^{-32}$ | $1.1 \times 10^{-14}$ | $9.5 \times 10^{-7}$ |
| M | Same-biology cross-batch agreement | n/a | 0.651 / 0.691 | 0.704 / 0.701 | 0.477 / 0.507 | 287.1 | $4.5 \times 10^{-63}$ | $9.4 \times 10^{-6}$ | $2.2 \times 10^{-20}$ | $4.5 \times 10^{-61}$ |
| M | Different-biology cross-batch agreement | n/a | 0.576 / 0.586 | 0.639 / 0.614 | 0.438 / 0.465 | 215.1 | $2.0 \times 10^{-47}$ | $9.7 \times 10^{-5}$ | $9.6 \times 10^{-14}$ | $3.2 \times 10^{-47}$ |
| M | Same- minus different-biology margin | n/a | 0.075 / 0.054 | 0.065 / 0.045 | 0.039 / 0.030 | 132.9 | $1.4 \times 10^{-29}$ | $1.9 \times 10^{-6}$ | $2.1 \times 10^{-30}$ | $2.5 \times 10^{-14}$ |
| N | LOBO3 macro F1 | all folds | 0.628 / 0.631 | 0.696 / 0.712 | 0.676 / 0.715 | 51.6 | $6.1 \times 10^{-12}$ | $2.7 \times 10^{-12}$ | $6.6 \times 10^{-7}$ | 0.0019 |
| N | LOBO3 macro F1 | multiclass folds | 0.645 / 0.657 | 0.666 / 0.678 | 0.664 / 0.678 | 11.8 | 0.0028 | 0.0014 | 0.0016 | 0.66 |
| N | LOBO3 macro AUC | multiclass folds (only) | 0.878 / 0.883 | 0.874 / 0.888 | 0.877 / 0.890 | 0.7 | 0.70 | 0.43 | 0.51 | 0.73 |
| N | LOBO2 macro F1 | all folds | 0.667 / 0.654 | 0.711 / 0.715 | 0.705 / 0.705 | 32.6 | $8.2 \times 10^{-8}$ | $3.9 \times 10^{-8}$ | $1.5 \times 10^{-6}$ | 0.17 |
| N | LOBO2 macro F1 | multiclass folds | 0.706 / 0.726 | 0.724 / 0.742 | 0.727 / 0.746 | 17.9 | $1.3 \times 10^{-4}$ | $3.3 \times 10^{-4}$ | $1.9 \times 10^{-5}$ | 0.60 |
| N | LOBO2 macro AUC | multiclass folds (only) | 0.888 / 0.901 | 0.879 / 0.904 | 0.885 / 0.904 | 1.5 | 0.48 | 0.91 | 0.55 | 0.22 |

In contrast, high-harshness methods lowered both the same-biology (Supplementary Figure 13C) and the different-biology cross-batch agreement (Supplementary Figure 13D). Also, the high harshness approaches had the smallest same-minus-different biology agreement margin (Supplementary Figure 13E) and the smallest margin gain over the unharmonized expression baseline (Supplementary Figure 13F). These findings suggest that reshaping samples into a common distribution by high harshness harmonization corrupted the rank structure shared by different batches of the same biology together with the gene expression structure that distinguishes batches of different biology.

On LOBO prediction the harshness order reversed: the low tier had the lowest three-class macro F1 on all folds (Supplementary Figure 13G) and on multiclass folds (Supplementary Figure 13H), and the lowest two-class macro F1 on both all class and multiclass only cohorts (Supplementary Figure 13J, K, respectively), while the medium and high tiers were close to each other on multiclass cohorts (Table 1). This reversal pattern can be explained by the fact that methods applying a per-batch monotone transform (low harshness) presumably keep the batch structure that a classifier trained on the other batches cannot predict, so a held-out batch remains displaced from the training batches. Neither the three-class nor the two-class AUC separated the tiers (Supplementary Figure 13I, L; Table 1). The permutation background was situated below every harshness tier, with medians of 0.237 to 0.261 for the three-class F1 and 0.500 for both AUCs as expected (Supplementary Figure 13G, I, J, L, respectively).

Taken together, we showed that prediction quality metrics (class N) are better in medium and high harshness approaches, whereas cross batch agreement by biomarker genes decreases in high harshness. Across all approaches, the class L biomarker correlation weakly decreased as the all cohorts three-class LOBO F1 increased due to low biomarkers correlation in 10_mnn and 12_scanorama methods (Spearman ρ = −0.234, p = 3.2 × 10^-29^, Supplementary Figure 14A, B by methods and strategies, respectively). The class M margin between same biology and different biology, in contrast, increased with the same F1 metric but again in a weak, noisy manner (ρ = 0.358, p = 2.3 × 10^-68^, Supplementary Figure 14C, D by methods and strategies, respectively). The cross-batch agreement margin gain over the unharmonized baseline increased very weakly with the multiclass-only F1 (ρ = 0.127, p = 2.6 × 10^-9^, Supplementary Figure 14E, F by methods and strategies, respectively). These opposite trends could indicate a tradeoff between biology subtype prediction and biomarker correlation preservation metrics, and in the cross-correlation clustermap of 325 metrics the class L, M and N metrics each form a separate block, with group N tending to be closer to global PCA-based metrics (Supplementary Figure 15; the 55 pairwise correlations are given in Supplementary File 1, sheet Scatter_correlations). A harmonization benchmark that aims at downstream prediction should therefore report class N next to classes L and M, because selecting on marker correlation alone could favor the approaches that predict held-out batches with poorer performance.

Moreover, the fact that high harshness methods performed worse by local, global and biology correlation metrics (classes L and M) but showed superior performance by distributional convergence and biology classes prediction suggests that these methods may get overfitted during harmonization. Indeed, the methods had biology labels as input arguments during harmonization (the column Diagnosis_cell_type_unified, described in (Nikitin et al. 2026 Sep 17)) and then they were assessed on LOBO prediction of these biology labels (with 2 or 3 classes but the larger class column Major_group)

### Preservation of biomarker expression and of cross-batch rank structure selects a distinct set of approaches

#### Class L: correlation of biomarker genes before and after harmonization

Since none of the 87 scoring metrics in the published benchmark assess expression of biomarker genes, we deeply studied performance of L and M metric classes in this study. These classes were visualized for the clustermap best approach in the benchmarking article (Nikitin et al. 2026 Sep 17) but they were not extensively compared between batch removal strategies and harmonization methods. We first plotted harmonized against unharmonized expression of the B cell lymphoma marker genes BCL6 and IRF4 and of the housekeeping gene EEF1A1 in the four largest cohorts, for four linear and four non-linear methods under the no-removal strategy with KNN imputation and without post-removal (Supplementary Figure 16A–H). The four linear methods, 03_limma, 05_combat, 29_combat_ref and 13_fsmvn, transformed every cohort along its own straight line, shifting and rescaling it, so the within-cohort order of the samples was kept exactly the same (Supplementary Figure 16A–D). Of the non-linear methods, 16_fsqn_r curved the cohort lines while preserving their order (Supplementary Figure 16E), and 17_quantile curved them and added noise around the curve, which lowered the within-cohort rank correlation (Supplementary Figure 16F). 10_mnn and 12_scanorama returned an almost constant value for each gene in every cohort, so the harmonized expression no longer followed the unharmonized one (Supplementary Figure 16G, H, respectively). These differences between methods were captured by the L and M class metrics. Class L evaluated the within-cohort order, which every linear method preserves by construction, and class M assessed the cross-batch rank structure, which was made identical across batches by collapse onto a constant profile.

We next examined which approaches can be selected by class L separately. Five methods including the 01_raw ‘method’ returned a median within-cohort marker correlation of exactly 1.000 (01_raw, 36_explobatch, 31_ruv3prps, 27_dwd and 21_harmonizr), and the low-harshness tier had a median of 0.998 (Table 1). Class L therefore favored output that is unchanged or close to unchanged: of its 112 best approaches, 55 are 01_raw and 38 are 36_explobatch. Seven approaches belonged to 13_fsmvn and 02_median_scaling, four approaches to 04_sva and one to 03_limma (Supplementary File 1, sheet Method_census_by_class). Because a marker correlation near 1.000 is reached by no transformation of the data, we used a marker correlation above 0.75 as a necessary condition, a quality control step that an approach must pass before the other classes are applied. We did not use the L class metrics as a ranking criterion.

#### Class M: cross-batch (dis)agreement of biomarker gene expression

We then examined the approaches selected using class M. Class M composite (average of all the class M metrics) was dominated by same-biology agreement, which increases whenever an algorithm transforms all expression profiles to a same profile, so class M elected only two methods: 10_mnn was in 61 of the top 112 places and 12_scanorama occupied the remaining 51. Both methods collapsed each biomarker gene onto an almost constant value as we showed previously (Supplementary Figure 16G, H), which maximizes same-biology agreement and removes the differences between biological groups at the same time.

#### Joint selection based on the L and M metrics

We then applied the two classes together as a joint gate, requiring a marker correlation above 0.75 and a positive margin of the same versus different biology cross-batch agreement. Of the 2,234 approaches, 1,686 (75.5%) passed it (Figure 4A, B). Across all approaches the margin increased weakly with the marker correlation (Spearman ρ = 0.177, p = 3.2 × 10^-17^), with the methods separating along both axes (Figure 4A) and the strategies mixed within the metrics-passing region (Figure 4B). The methods that passed with highest frequency were the linear ones with low or medium harshness: 06_combat_seq, 03_limma, 07_pycombat, 08_inmoose_combatseq, the ‘method’ 01_raw, 02_median_scaling and 05_combat. By strategy, the RNA-seq-only, FF-only and no-removal strategies passed most often (141, 130 and 130 approaches). The FFPE-only strategy passed least often, in 76 of its 164 approaches, presumably due to its high batch heterogeneity. Interestingly, the unharmonized output 01_raw passed the joint control in 80 of its 84 approaches, and this was more often than the other approaches (74.7%; Fisher’s exact p = 1.7 × 10^-6^), so passing the thresholds by L and M metrics does not require harmonization.

**Figure 4.**
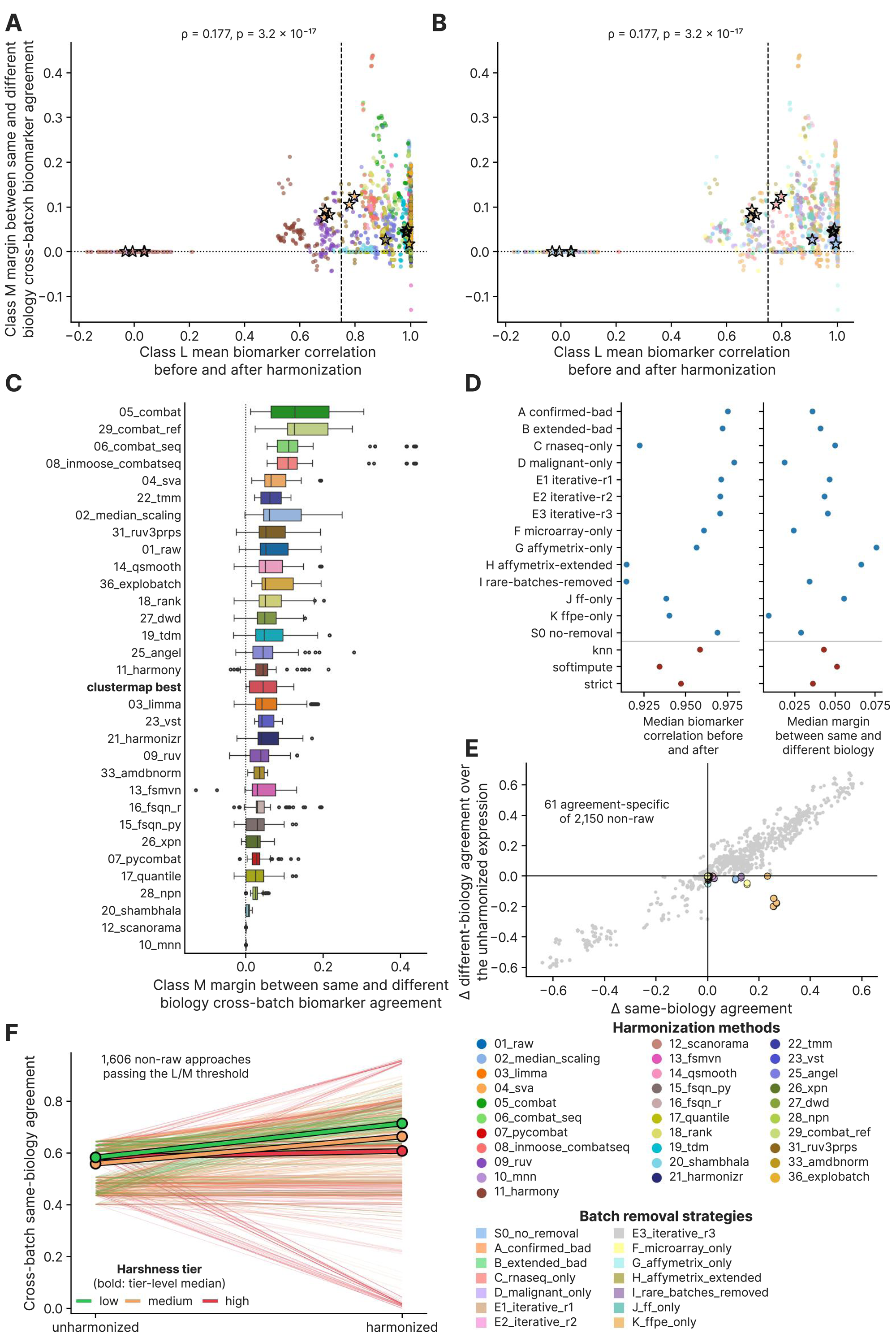
Marker-gene preservation and cross-batch rank structure, metric classes L and. **M.** (A) Scatterplot of the class M margin (same-biology minus different-biology cross-batch rank agreement) against the class L mean within-cohort marker correlation before and after harmonization for all 2,234 approaches, color annotated by harmonization method. The dashed vertical line is the 0.75 gate of class L and the dotted horizontal line is a margin of zero. (B) The same scatterplot color annotated by batch removal strategy. (C) Boxplots of the class M margin by harmonization method, with methods ordered by median margin; the 15 clustermap best approaches are pooled into one additional row, “clustermap best”, shown in bold. The dotted vertical line is a margin of zero. (D) Median class L marker correlation (left) and median class M margin (right) for each batch removal strategy (blue) and each imputation (red), each median taken over all approaches of that strategy or imputation. (E) Change in different-biology agreement against change in same-biology agreement, both taken against the unharmonized baseline 01_raw at the same strategy and imputation, for the 2,150 non-01_raw approaches. The 61 agreement-specific approaches, which raise same-biology agreement and lower different-biology agreement against their baseline, are situated in the lower right quadrant and are color annotated by batch removal strategy (ρ = 0.931, p < 10^-300^ over all points). (F) Same-biology agreement of the unharmonized baseline and of the harmonized output, one thin line per approach, for the 1,606 approaches other than 01_raw that pass the joint quality control (marker correlation above 0.75 and margin above 0). Lines are color annotated by method harshness tier, and the bold line of each tier joins the tier medians. The clustermap best approaches are shown as empty black stars in panels A and B. In panel C, boxes span the interquartile range (IQR) with the median marked, whiskers extend to 1.5 × IQR and points beyond them are drawn individually.

We then additionally compared the 1,606 L and M class passing approaches other than 01_raw with their unharmonized baseline. In these approaches, harmonization raised the median same-biology agreement over the unharmonized baseline by 0.113 in the low harshness tier and 0.110 in the medium tier and left it unchanged in the high tier (Kruskal–Wallis p = 1.1 × 10^-29^; Figure 4F). The high tier also contained most of the approaches that lost more than 0.1 of their cross-batch agreement after harmonization: 100 of its 685. The low tier contained no such approaches (Fisher’s exact p = 4.4 × 10^-25^).

#### Comparison of methods, strategies and imputations by the L and M metrics

We then compared batch-removal strategies, imputations and methods on the class M and L metrics (Figure 4C, D, respectively). Strategies differed in median marker correlation (Kruskal-Wallis p = 4.1 × 10^-4^), from 0.979 under malignant-only down to 0.914 under rare-batches-removed (Figure 4D). Strategies also had significant differences in median margin (p = 3.6 × 10^-33^), from 0.076 under Affymetrix-only down to 0.009 under FFPE-only (Figure 4D). Only two strategies increased the margin above their own unharmonized baseline in the median, and FFPE-only had the highest increase, by +0.018 (Wilcoxon signed-rank p = 1.2 × 10^-13^). RNA-seq-only raised the margin over the unharmonized dataset by +0.008 (p = 2.5 × 10^-4^). Eight strategies lowered the margin below their baseline (p ≤ 6.5 × 10^-9^ each), and the remaining four strategies changed it by less than 0.001 in the median (Supplementary File 1, sheet Paired_tests). The criterion of the margin corrected by the unharmonized matrix can be considered as the most stringent for the class M metrics, since it requires that a harmonization method amplifies biology differences that are initially weak compared to the batch-related differences.

Imputations did not separate the median marker correlation (Kruskal-Wallis p = 0.13, Figure 4D), but they had significant differences by the median margin (p = 4.4 × 10^-5^). The median margin was 0.051 under softimpute, 0.043 under KNN imputation and 0.036 under strict exclusion. Harmonization methods had the most significant separation: median marker correlation changed from 1.000 for five methods to −0.008 for 12_scanorama (Kruskal-Wallis p < 10^-300^; Supplementary File 1, sheet Paired_tests). Median margin changed from 0.126 for 05_combat to 0.000 for 10_mnn and 12_scanorama (p = 1.9 × 10^-194^, Figure 4C).

#### Increasing same biology agreement and different biology disagreement leads to a group of agreement-specific approaches

We then additionally required the two directions of class M separately: an acceptable approach had to increase same-biology agreement and decrease different-biology agreement against its unharmonized baseline. Only 61 of the 2,150 non-raw approaches (2.8%) adhered to this criterion, and we termed them agreement-specific (Figure 4E). This low percentage can be explained by the fact that the two changes were strongly correlated across approaches (Spearman ρ = 0.931, p < 10^-300^), i.e. most approaches moved both agreements in the same direction.

The agreement-specific approaches concentrated in the FFPE-only strategy, with 21 approaches comprising 13.3% of its non-raw approaches against 2.0% in the other strategies (Fisher’s exact p = 4.0 × 10^-10^). The microarray-only strategy followed with 11 agreement-specific approaches (p = 2.6 × 10^-3^). The seven approaches of the rare-batches-removed strategy did not exceed its fraction in the total benchmark (p = 0.19).

Imputations did not show an enrichment in the agreement-specific approaches (Fisher’s exact p ≥ 0.27 for each imputation against the other two). Among the harmonization methods, 05_combat contributed 14 approaches (Fisher’s exact p = 1.7 × 10^-8^), and the next methods were 36_explobatch, 06_combat_seq and 08_inmoose_combatseq, with six approaches per method (p ≤ 0.028). The full list of the 61 approaches is provided in Supplementary File 2, sheet Agreement_specific_61.

#### The two winners: ComBat-Seq in FFPE-only under KNN imputation

The two approaches highest by margin gain were two different methods of the same strategy and imputation: FFPE-only with KNN imputation and no post-removal, under 06_combat_seq and under 08_inmoose_combatseq. Both approaches had a margin of 0.439 and a gain of +0.457 over their own unharmonized baseline. The gain of margin consisted of +0.257 in same-biology agreement and −0.200 in different-biology agreement. These two implementations of ComBat-Seq agreed to six decimals in their metrics, which could be utulized as a positive control on the class M computation.

We then checked the expression output scale of the two methods behind these best margins. 06_combat_seq and 08_inmoose_combatseq are also the two methods whose output maxima exceeded their 99th percentiles by seven or more orders of magnitude (Figure 3E). Classes L and M are built on rank correlations, which do not change under the output rescaling and cannot detect this defect. Therefore, a separate controplof the output scale is needed to supplement these metrics.

Taken together, the analysis of classes L and M showed that marker correlation saturates for the methods that change expression least, and that cross-batch agreement is highest for the methods that collapse expression onto a constant profile, so neither class can select the best approaches alone. A harmonization benchmarking pipeline should use marker correlation as a initial quality gate, read cross-batch agreement through the same-minus-different-biology margin relative to the unharmonized baseline, and check the scale of the harmonized output separately.

### Cross-batch prediction results in a third set of best approaches, led by the ComBat family

#### Single class batches were predicted with higher performance than the multiclass batches

We then evaluated class N metrics, which evaluate a classifier fitted on all the batches expect a held-out one, to predict the biology classes of this held-out batch. Over the 2,234 approaches, this leave-one-batch-out (LOBO) prediction of the three-class target produced 32,746 held-out folds over 24 RNA batches. Of these folds, 15,436 (47.1%) contained a single biological class, and other folds carried two or three classes. Single-class folds reached a median macro F1 of 0.903, against 0.733 for the multiclass folds (Mann–Whitney U, p = 2.3 × 10^-221^; Figure 5C). Among the multiclass folds, two-class folds performed poorer than three-class folds (median 0.629 against 0.804, Mann–Whitney U, p < 10^-300^; Figure 5C).

**Figure 5.**
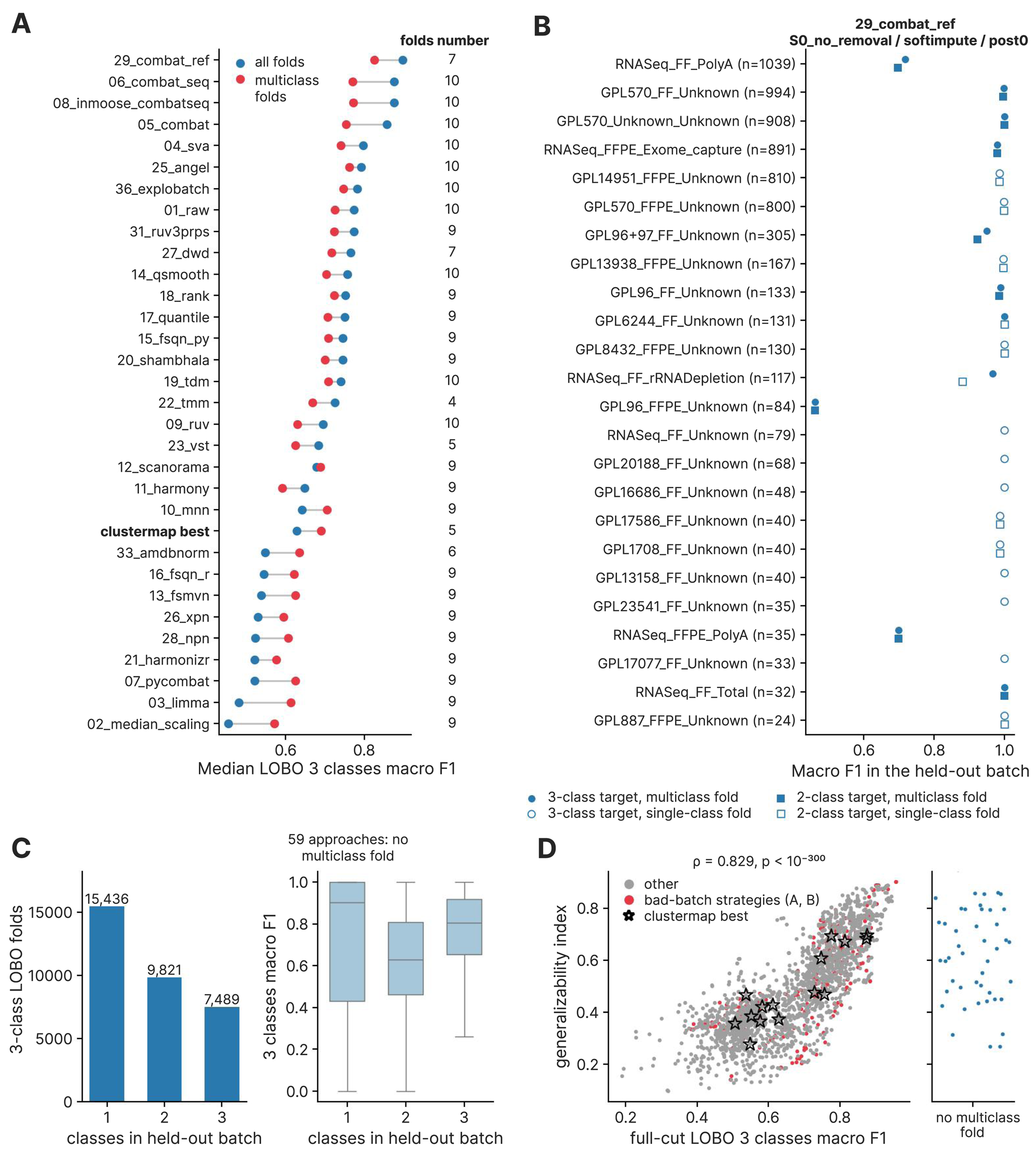
Cross-batch prediction, metric class. **N.** (A) Median leave-one-batch-out (LOBO) three-class macro F1 by harmonization method, over all held-out batches (blue) and over the multiclass folds only (red), with methods ordered by the all-fold median three-class macro F1; the 15 clustermap best approaches are shown one additional row in bold. The column on the right gives the median number of multiclass folds per approach of each method. (B) Macro F1 in each held-out batch for S0_no_removal__softimpute__29_combat_ref__post0, the eligible approach of the no-removal strategy with the highest full-cut three-class macro F1, for three-class batches (circles) and two-class batches (squares). Batches are ordered by sample count, given in brackets, and hollow markers are single-class folds: 13 of the 24 three-class folds and 9 of the 18 two-class folds carried one biology class. (C) Fold composition by classes number per batch. Left subpanel: number of three-class LOBO folds by the number of biology classes in the held-out batch, with the count above each bar. Right: macro F1 of the same folds by the number of classes, drawn as boxplots without outlier points; the title gives the 59 approaches that have no multiclass fold. (D) Generalizability index against the full-cut LOBO three-class macro F1 for the approaches with at least one multiclass fold. The generalizability index is the mean of six rank percentiles: 1) the full-cut and 2) multiclass-only three-class macro F1, 3) the full-cut two-class macro F1, 4) the class L mean marker correlation, 5) the class M margin gain over the unharmonized baseline, and 6) the worst multiclass fold (the lowest three-class macro F1 over the multiclass folds of the approach). Red points are the two strategies that are based on a visual assessment of batches (A_confirmed_bad and B_extended_bad), gray points are all other approaches, and empty black stars are the clustermap best approaches. The narrow panel on the right shows the generalizability index of the 59 approaches with no multiclass fold, jittered horizontally, since the full-cut LOBO F1 score is not defined for these approaches.

We also examined why held-out batches with a single class had higher F1 scores. If only one class is present, the macro F1 of the fold reduces to the F1 of that class, and the fold tests whether the classifier assigns the diagnosis of the whole held-out batch and does not test whether it separates diagnoses. Surprisingly, these two fold types (single and multiclass) had close means (0.690 against 0.679) although their medians differed, so the difference was connected to the shape of the distribution. Single-class batches separated into the batches that labeled correctly in the median fold, such as several FF microarray batches at 1.000, and those that failed, such as the FFPE batches GPL887_FFPE_Unknown at 0.000 and GPL13938_FFPE_Unknown at 0.069 (Supplementary File 2, sheet Fold_composition). A mean over all folds therefore mixed the assignment of whole cohorts with the separation of diagnoses measured on the multiclass folds. For this reason, we analyzed class N values on two settings, or cuts, simultaneously: the full cut (the mean over all held-out batches) and the multiclass-only cut (the mean over the folds that carry at least two classes).

#### Selection of the best approaches by prediction quality

We then compared the two cuts across approaches. The full and the multiclass-only means correlated with Spearman ρ = 0.810 over the 2,174 approaches containing both cuts (p < 10^-300^). Over the 2,058 approaches with at least three multiclass folds the correlation was ρ = 0.815 (p < 10^-300^). This agreement was high enough for both cuts to place the same method, 29_combat_ref in prediction best approaches (Figure 5A), so we selected the best approaches based on the both cuts performance (Table 2).

**Table 2.**
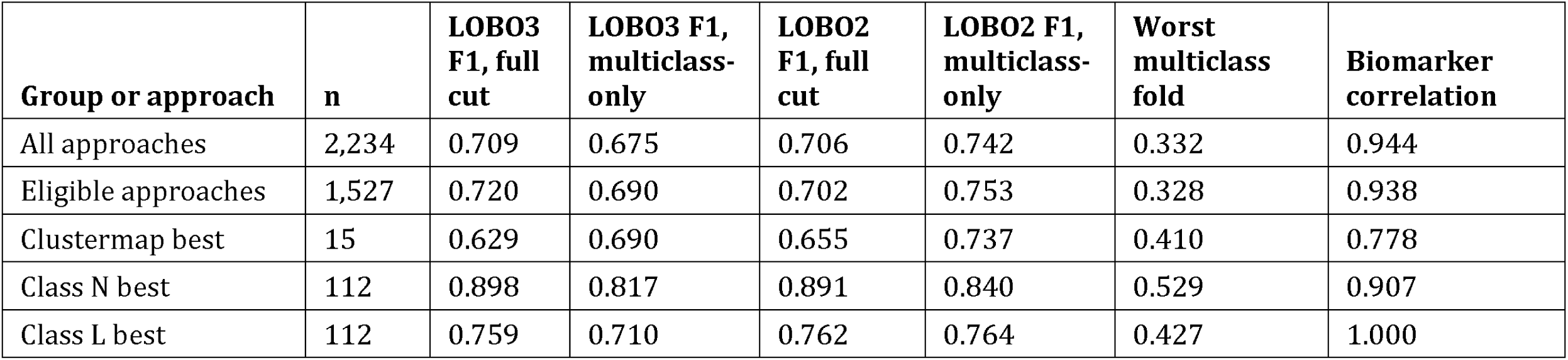
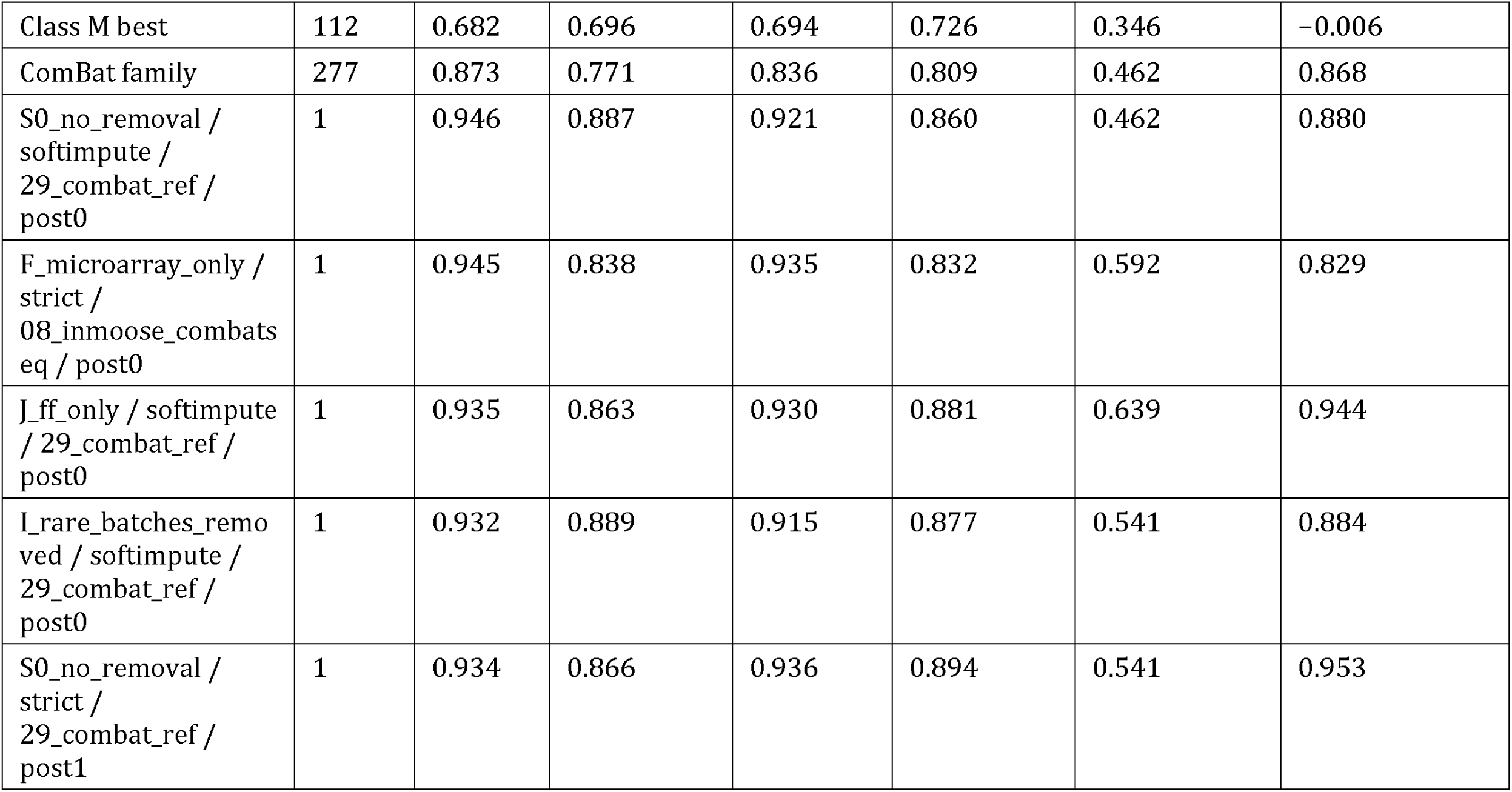
Leave-one-batch-out prediction metrics for the selected groups and the leading approaches. LOBO3 is the three-class target (DLBCL, FL and normal B cells); LOBO2 is the two-class target (FL against DLBC. The worst multiclass fold is the lowest three-class macro F1 over an approach’s multiclass folds. Marker correlation is the class L mean within-cohort marker correlation before and after harmonization. Group rows contain medians over the approaches of the group. The best eligible approach is given for the four strategies with the highest full-cut LOBO3 F1. Detailed metrics for this table can be found in Supplementary File 2. ComBat family methods is 05_combat, 06_combat_seq, 08_inmoose_combatseq and 29_combat_ref.

We next identified the fold structures with high prediction quality value carrying no information on class separation. All 59 approaches without a multiclass fold belonged to the Affymetrix-only strategy with post-removal, and 20 of them showed a full-cut F1 of at least 0.999. We excluded these approaches from the best ones by prediction quality (Table 2).

For reliable selection of the best approaches, we defined a separate eligible set by excluding the two strategies that removed batches by visual assessment in the previous benchmark (Nikitin et al. 2026 Sep 17), A_confirmed_bad and B_extended_bad, and the approaches with fewer than five multiclass folds. This eligible set contained 1,527 approaches. The best eligible approach of each of the four best strategies used 29_combat_ref, except under the microarray-only strategy, where 08_inmoose_combatseq was the top method (Table 2).

The absolute best approach, no removal with softimpute under 29_combat_ref without post-removal, reached a full-cut F1 score of 0.946 and a multiclass-only F1 mean of 0.887 over 11 multiclass folds. We visualized per-batch performance by F1 score for two and three class prediction, for all and multiclass-only batches (Figure 5B). Of the 24 total folds, 13 were single-class and had F1 scores between 0.986 and 1.000. The poorest performance we found for GPL96_FFPE_Unknown, an FFPE microarray batch of 84 samples and two classes (F1 0.462, AUC 0.543). The next two batches by the worst performance were the RNA-seq batches RNASeq_FFPE_PolyA (F1 0.698) and RNASeq_FF_PolyA (F1 0.718; Supplementary File 2, sheet Per_batch_folds). Two of these three hardest folds were FFPE batches, presumably because degraded FFPE material shifts expression profiles away from those of the training batches.

#### ComBat family is the best by prediction quality

We deeply examined the 112 best approaches by class N. These approaches mostly belonged to the ComBat family: four implementations of that algorithm, 05_combat, 29_combat_ref, 06_combat_seq and 08_inmoose_combatseq, occupied 84 of the 112 places, and eight further methods shared the remaining approaches (Supplementary File 1, sheet Method_census_by_class). It should be noted that in the total approaches set the ComBat family methods reached a median full-cut F1 of 0.873 against 0.669 for the other methods (Mann–Whitney U, p = 2.9 × 10^-105^). The same difference persisted on the multiclass-only folds (0.771 against 0.665, Mann–Whitney U, p = 1.0 × 10^-44^).

Harmonization methods in general significantly differed between each other on the full cut of LOBO 3 classes F1 score (Kruskal–Wallis H = 1,425, p = 4.3 × 10^-281^; Figure 5A). Their medians changed from 0.898 for 29_combat_ref to 0.453 for 02_median_scaling, and the same two methods were first and last by the multiclass-only cut (0.827 against 0.571, Kruskal–Wallis p = 3.2 × 10⁻¹⁶⁹). The five poorest methods by the full cut, 02_median_scaling, 03_limma, 07_pycombat, 21_harmonizr and 28_npn, include four of the six methods that increase batch PCReg to saturation (Figure 5A). Therefore, these methods can be interpreted as removing the diagnosis-associated variance together with the batch-associated one. 07_pycombat, a Python implementation of ComBat, is one of these four approaches and was not included into the class N best set (Supplementary File 1, sheet Method_census_by_class).

The class N best set contained 13 of the 14 strategies, and the FFPE-only strategy was not included (Fisher exact test, p = 2.7 × 10^-4^). By imputations we observed a weak difference from the total approaches set of imputations (χ^2^ goodness-of-fit test, p = 0.042): there were 50 approaches with strict exclusion, 33 with softimpute and 29 with KNN imputation. By post-removals there were no significant differences (63 approaches with post-removal, two-sided binomial test, p = 0.22).

#### Class N prediction quality metrics comparison by methods, strategies and imputations

We then compared the pipeline factors (methods, strategies, imputations) across all approaches by LOBO 3 classes F1 score on all and multiclass-only cohorts (Supplementary File 1, sheet Paired_tests). Strategies separated the full-cut F1 significantly (Kruskal–Wallis p = 1.5 × 10^-17^), with group medians from 0.763 for Affymetrix-extended to 0.561 for FFPE-only, whereas imputations separated it less significantly (Kruskal–Wallis p = 1.7 × 10^-4^). The effect sizes order the three factors favored harmonization methods: ε² was 0.634 across methods, 0.044 across strategies and 0.007 across imputations.

In the case of the multiclass-only cut, strategy medians changed from 0.724 for microarray-only to 0.462 for Affymetrix-only (Kruskal–Wallis p = 3.7 × 10^-80^). The Affymetrix-only median was calculated on the 95 of its 154 approaches that carried multiclass batches. Importantly, post-removal increased the full-cut median from 0.692 to 0.719 (Mann–Whitney U, p = 2.9 × 10^-3^) but did not change the multiclass-only median (0.669 against 0.681, Mann–Whitney U, p = 0.13), which indicates that post-removal imrpved performance for single-class folds only.

#### Random background control of the class N prediction quality metrics

In the original harmonization benchmark, we controlled every class N value with 100 permutations of the biological labels (Nikitin et al. 2026 Sep 17). Under permutation, the full-cut three-class macro F1 decreased to the harshness tier medians between 0.237 and 0.261, against an observed median of 0.709 over all approaches (Supplementary Figure 13G). The permuted two-class F1 dropped to the harshness tier medians between 0.393 and 0.441 (Supplementary Figure 13J). The permuted AUC stayed at 0.500 for both targets in every tier, the value expected from a random classifier (Supplementary Figure 13I, L). The observed full-cut three-class F1 exceeded its permutation distribution at p ≤ 0.05 in 2,197 of the 2,233 approaches carrying the column, confirming that the observed prediction performance was not random.

#### Generalizability index composed of the correlation and prediction quality metrics

We then combined six prediction and preservation columns into the generalizability index, the mean of six rank percentiles: 1) the full-cut and 2) multiclass-only three-class macro F1, 3) the full-cut two-class macro F1, 4) the class L marker correlation, 5) the class M margin gain over the unharmonized expression matrix and 6) the worst multiclass fold by 3 classes LOBO F1 score. Over all the approaches, the 15 highest index values, from 0.949 to 0.986, belonged to Affymetrix-only approaches with post-removal and without multiclass folds. We excluded these approaches, since their high generalizability index was artificially inflated by single class only high F1 scores (Figure 5D). Under restriction to the 1,527 eligible approaches, the maximum index was observed in approach of no removal with strict exclusion under 29_combat_ref with post-removal at 0.886 (Table 3). All 15 leading eligible approaches were ComBat implementations, nine under 29_combat_ref and six under 05_combat, and nine of the 15 used the FF-only strategy. The generalizability index followed the full-cut three-class F1 (Spearman ρ = 0.829, p < 10^-300^; Figure 5D). The 15 clustermap-selected approaches had ranks between 433rd and 2,022nd of the 2,234 total approaches, and their median generalizability index did not differ from that of the other approaches (0.466 against 0.476, Mann–Whitney U, p = 0.95).

**Table 3.**
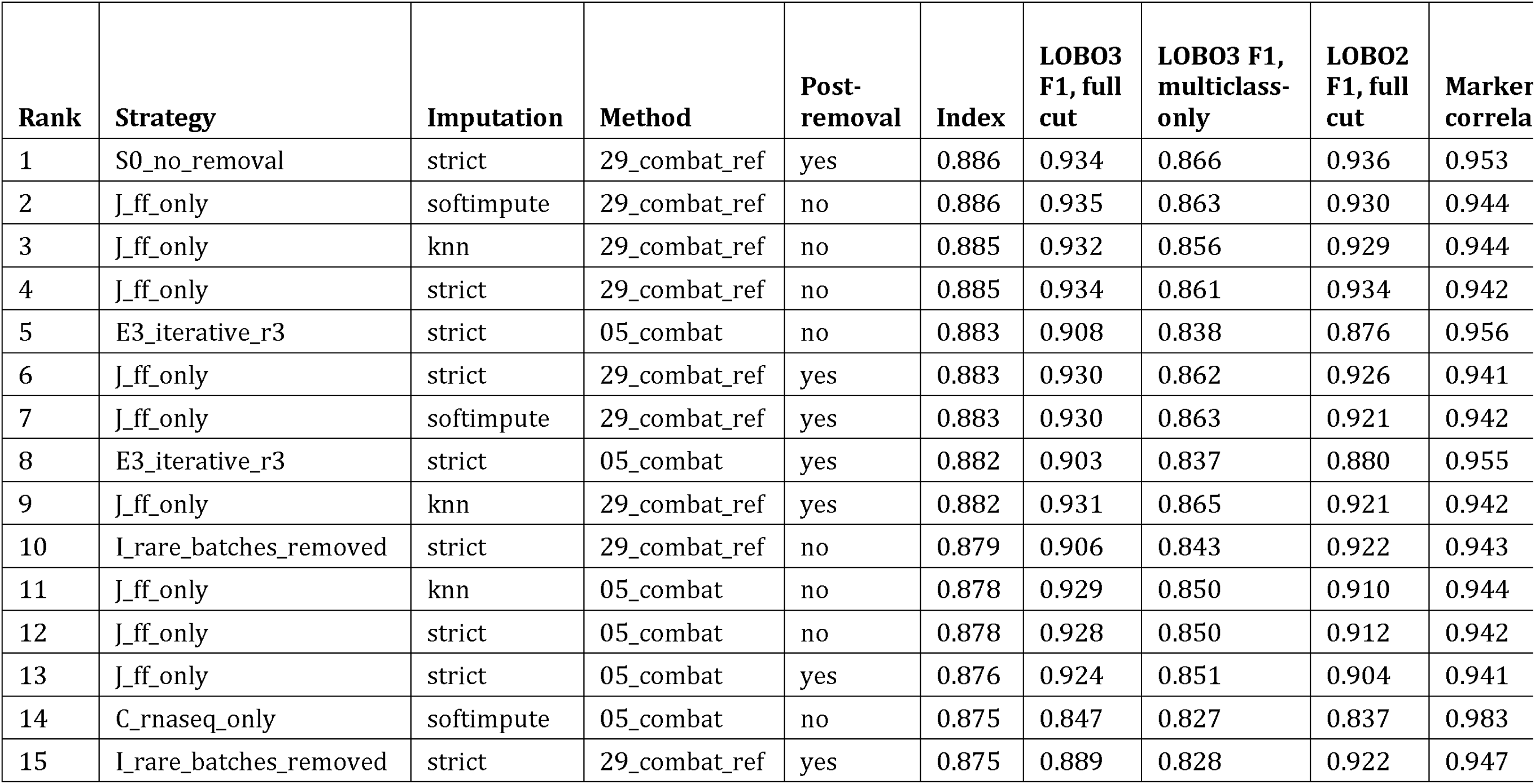
The 15 eligible approaches with the highest generalizability index and its six components. The table gives the raw value of each component. Eligible approaches exclude the two strategies A and B and approaches with fewer than five multiclass folds (n = 1,527). Ranks are among the eligible approaches. The full ranking of all 2,234 approaches is in Supplementary File 1, sheet Generalizability_ranking.

Taken together, cross-batch prediction resulted in a best approaches set dominated by the ComBat family, and the leading approaches preserved this composition on both fold cuts and under the generalizability index. Nearly half of the held-out batches contained a single diagnosis, so mean over all folds mized the assignment of whole cohorts with the separation of diagnoses, and strategies retaining batches confounded with diagnosis reached the highest values. A benchmark that uses held-out prediction should therefore report the multiclass-only cut and the multiclass fold count beside the full-cut values, exclude the strategies that retain confounded batches before ranking, and compare the observed values with a label-permutation random expectation.

### The best sets elected by the individual metric classes show limited intersection

#### Cross-comparison of the best approach sets

As a final step of harmonization quality metrics analysis and benchmarking, we selected the best approaches per each class over the same 2,234 approaches and compared the resulting eight sets of 112 approaches with each other and with the 15 approaches of the published clustermap selection (Figure 6A; Supplementary File 2, sheet Cross_election_pairs). Every overlap between the sets was tested against a random expectation using a one-sided hypergeometric test in the direction observed. The largest agreement was between the local class and the composite index, resulting in 71 shared approaches (Jaccard 0.464, p = 9.5 × 10^-75^), which is expected since local metrics are the most distinctive part of the composite index (Supplementary Figure 11C, D). The composite top approaches also shared 51 approaches with the structural metrics, 50 with the distributional class and 20 with the global class (Supplementary File 2, sheet Cross_election_pairs), again as expected. Among the four component classes, the local and distributional classes agreed most, with 51 shared approaches (p = 4.9 × 10^-41^). The other five pairs shared between 8 and 18 approaches. Of the ten pairs formed by the four component classes and the composite, only the global and local classes did not agree above chance (8 shared against 5.6 expected, p = 0.20), indicating orthogonality of local and global batch correction performance.

**Figure 6.**
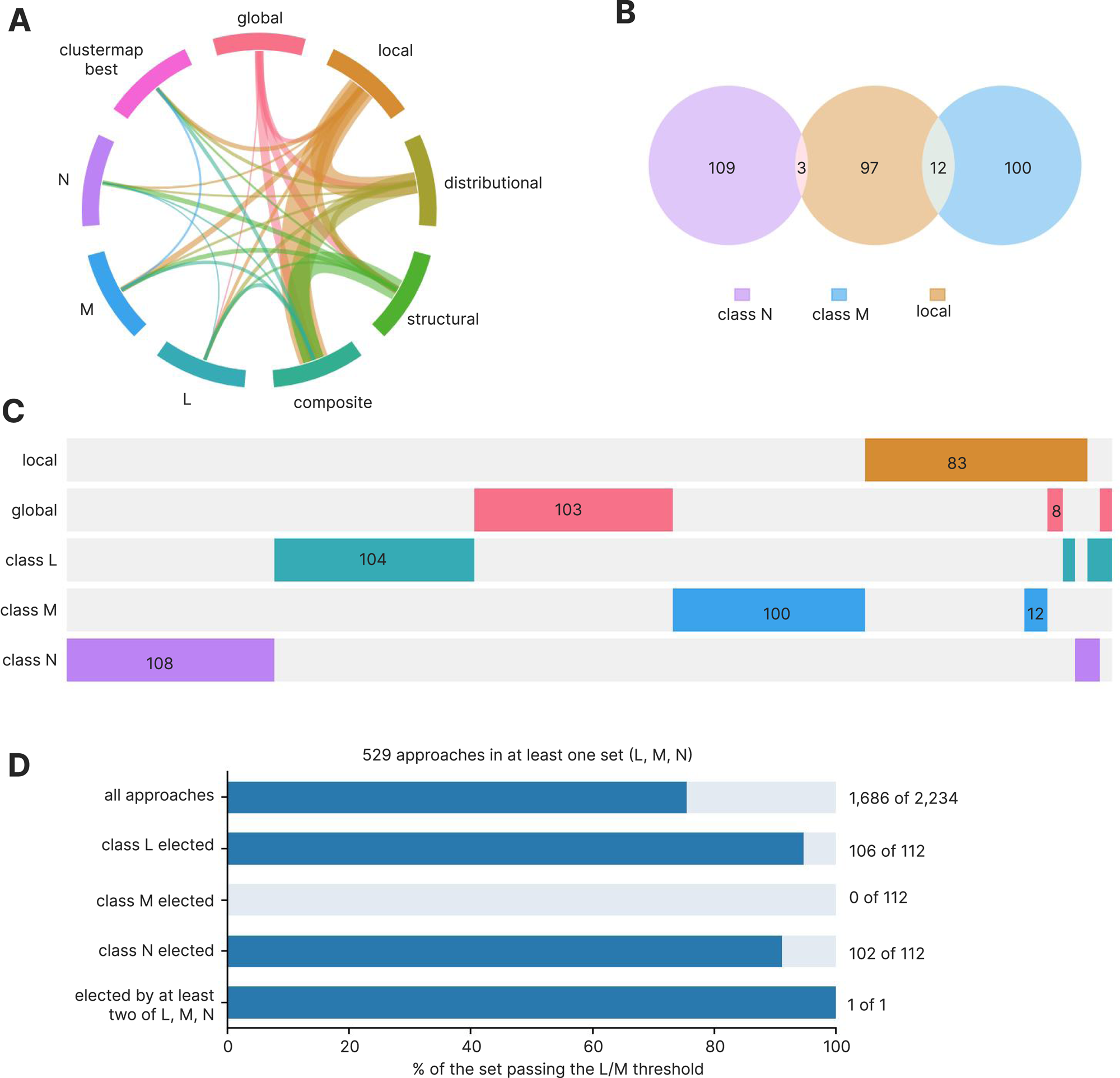
Overlap of the sets of best approaches elected by the individual metric classes. Each metric class resulted in the top 5% of the 2,234 approaches by its class score (mean rank percentile of its metrics), 112 approaches per class. (A) Chord diagram of the nine best approach sets: the global, local, distributional, structural and composite classes, classes L, M and N, and the 15 clustermap best approaches. A ribbon joins two sets that share at least one approach, its width grows with the number of shared approaches pair (counts in Supplementary File 2, sheet Cross_election_pairs). (B) Venn diagram of the class N, class M and local metrics best sets. (C) Chunk plot of the intersections between five sets of classes N, M and L and of the global and local classes. Each column is one membership pattern, colored cells mark the sets that contain it, and the column width is the number of approaches with that pattern; counts are shown under columns of at least 8 approaches. (D) Barplot showing fraction of approaches that pass the L/M quality control gate (marker correlation above 0.75 and margin above 0), the approaches range from all 2,234, the three sets elected by classes L, M and N, and the approaches elected by at least two of the three; bar height is proportional to the fraction. The single approach elected by two of the three classes is I_rare_batches_removed__softimpute__01_raw__post1, an unharmonized baseline elected by classes L and N, a trivial result.

We then compared the three biology classes with each other and with the remaining sets. Classes L and M shared no approach, and neither did classes M and N, against 5.6 expected by random background (hypergeometric p = 0.0027 for each pair). Classes L and N shared a single approach, which was also fewer than expected (p = 0.020). This approach was the unharmonized output of the rare-batches-removed strategy with softimpute and post-removal, which was a trivial result without technical or biological significance. Class N shared no approach with the global class (p = 0.0027), although class N tested prediction quality by logistic regression in top 10 principal components. It shared two approaches with the composite index, which was not below the random expectation (p = 0.071), and its largest overlap was 10 approaches with the structural class (Jaccard 0.047). Class M shared 12 approaches with the local class (p = 0.0088). Class L shared at most 7 approaches with any other class, and the only one of its overlaps that was significantly different from the random exprectation was the single approach shared with the global class (p = 0.020). We visualized a Venn diagram of class N, class M and local sets in Figure 6B and chunk intersection plot of L, M, N, local and global metrics in Figure 6C.

#### Comparison of the clustermap best approaches to the eight best approach sets

We then compared the 15 clustermap-selected approaches with the eight elected sets. They shared nine approaches with the local class (Jaccard 0.076, p = 5.7 × 10^-9^), 7 approaches with the composite and 6 with the distributional class, both significantly above the random expectation (Supplementary File 2, sheet Cross_election_pairs). Also, the clustermap approaches shared 3 approaches each with the structural class and class M (p = 0.036), and none with the global class, class L or class N. For a set of 15, sharing no approach with a random set of 112 had a probability of 0.46 (hypergeometric test), so the empty overlaps with classes L and N did not show significant disagreement. The clustermap group also did not differ from the other approaches on prediction quality: its median full-cut F1 of 0.629 against 0.709 was not a significant difference (Mann–Whitney U, p = 0.97; Table 2), and the same was observed for the generalizability index (p = 0.95).

#### Harmonization methods favored by the best approach sets

We then counted the methods inside each of the best elected sets (Supplementary File 1, sheet Method_census_by_class). The method distribution was highly non-uniform: four metric classes concentrated at least three quarters of their places in two to four methods, different for each metric class. The global class had 42 of its 112 places to 03_limma and 42 13_fsmvn. Both methods were among the six methods that increased batch PCReg to the saturation level above 0.999. Class L had 93 best approaches out of 112 being 01_raw and 36_explobatch, class M shared all the 112 approaches between 10_mnn and 12_scanorama, and class N gave 84 places to the four ComBat implementations. The local, distributional, structural and composite sets were more diverse, each of them spread over 20 to 25 methods. The distributional, structural and composite best approach sets were led by 16_fsqn_r with 12 to 18 places, whereas the local set was dominated by 10_mnn with 19 places. In total, of the 2,234 approaches, 1,686 passed the joint threshold of a marker correlation above 0.75 and a positive margin of cross-batch agreement of same versus the different biology classes (Figure 6D).

We then asked which methods each class would recommend if a benchmark analyzed that class alone. As we noted in the previous passage, the global class would recommend 03_limma and 13_fsmvn. 13_fsmvn had a median full-cut F1 of 0.537 against 0.715 for the other approaches which was significantly lower (Mann–Whitney U, p = 3.6 × 10^-18^), 03_limma was second to last of the 31 methods by this metric (Figure 5A). Class M, read without the margin, would recommend 10_mnn and 12_scanorama, the methods completely corrupting biology as we showed earlier. Class N would recommend the ComBat family, whose count-based implementations 06_combat_seq and 08_inmoose_combatseq returned mean maximum expression values of 2.76 × 10^8^ and 1.46 × 10^11^ (Figure 3E). Class L would recommend unharmonized or near-unharmonized output, 01_raw and 36_explobatch (Supplementary File 1, sheet Method_census_by_class). In each of the four cases, the recommended methods scored poorly on at least one quantity measured by another class.

#### Final benchmarking recipe for cross-platform transcriptomic bulk datasets harmonization

Taken together, the four batch-facing metric classes (global, local, distributional, structural) and the composite agreed above chance in nine of their ten pairs, whereas the three biology-facing classes shared fewer approaches than random expectation with each other and elected sets built on different methods (Supplementary File 2, sheet Cross_election_pairs). A benchmark that selects harmonization approaches should therefore explicitly describe its metric classes, its quality control thresholds and its held-out test before ranking, and report the best approaches set of every class with the intersections and their chance expectation. Class L metrics of biomarkers correlation before and after harmonization should serve as a gate to select approaches that preserve biomarker expression rank structure (Table 1). Class M metrics of cross-batch agreement should be defined as the margin of same biology versus the different, corrected against the unharmonized baseline at the same strategy and imputation, because agreement alone rewards algorithms that converge all expression profiles regardless of their biology. Class N prediction quality metrics should be reported on both fold cuts with the multiclass fold count, because in the dataset analyzed 47.1% of the held-out batches contained one biology class.

Based on these insights, we build a final benchmarking recipe for cross-platform transcriptomic bulk datasets harmonization (Figure 7). We propose to keep the following metrics in a generalizable benchmarking: PCReg, LISI, kBET, tSNE/UMAP entropy, LOBO macro F1 on single class and multi class batches, correlation of biomarker genes before and after harmonization and cross-batch agreement of biomarker genes between the same versus the different biology. The proposed recipe and the metrics can be implemented using the ComboBatch tool we released in the previous benchmark publication, available via the link https://github.com/Nikit357/ComboBatch.

**Figure 7.**
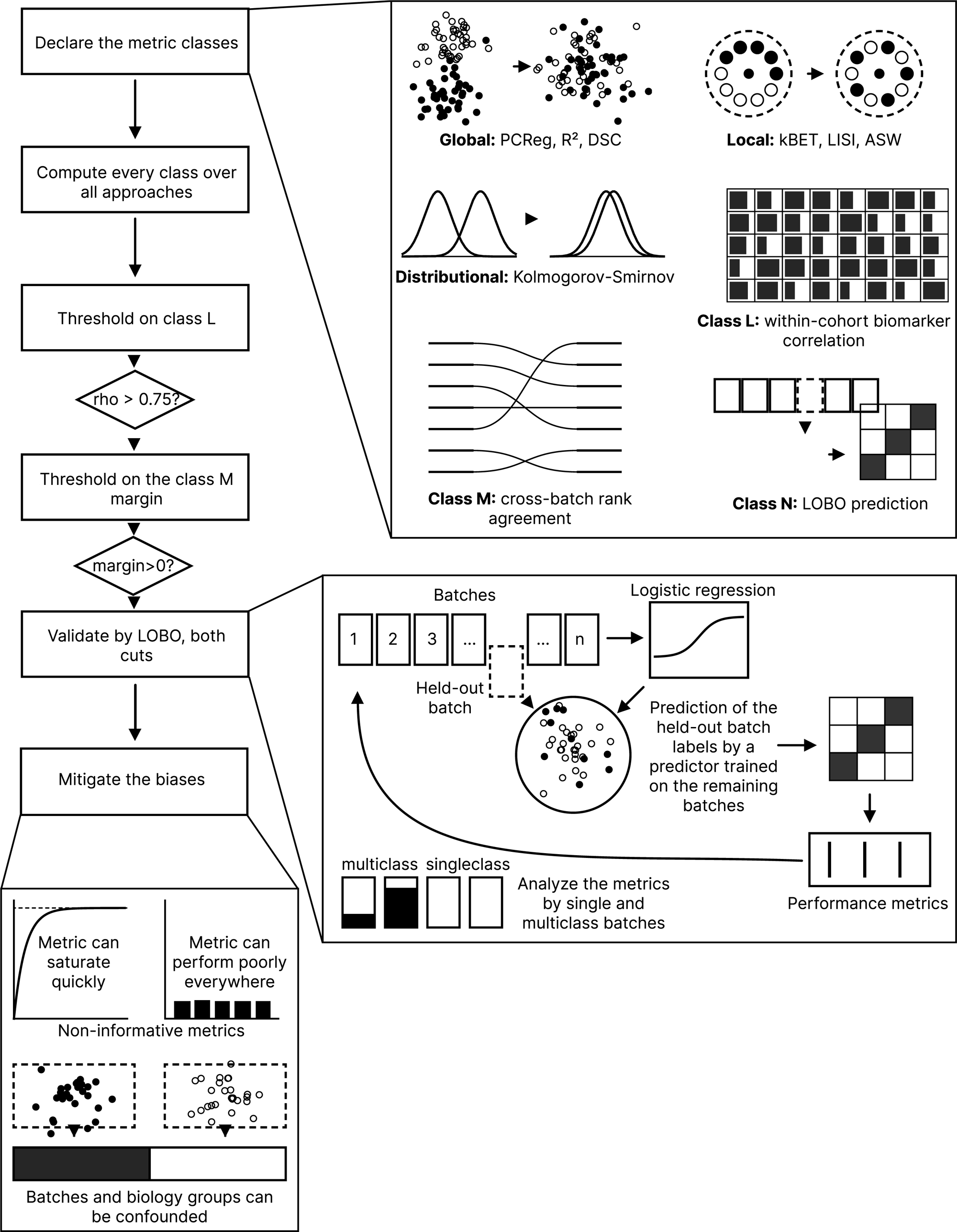
A recipe for selecting metrics when benchmarking harmonization approaches. A block scheme in six steps that are required for a generalizable benchmark for harmonization approaches. The three insets describe in a greater detail: 1) metrics classes, 2) leave-one-batch-out (LOBO) prediction quality metrics and 3) benchmarking biases.

### Conclusions

Selecting the best harmonization approach is essential for cancer biomarker discovery and validation in bulk transcriptomics. We aimed to develop a generalizable pipeline of this selection using the set of 2,234 harmonization approaches with 31 methods and 3 imputations, each approach evaluated using 334 quality metrics (Nikitin et al. 2026 Sep 17).

Based on eight metric classes, we extracted the top 5% of 2,234 harmonization approaches for each class and compared the elected sets with the 15 approaches selected by the published clustermap (Nikitin et al. 2026 Sep 17). Best sets of the biology preserving and prediction quality metrics (classes L, M, N) overlapped significantly below the random background between each other and with the global, local, and distributional metrics best sets.

Compared to the FSQN and SVA methods showed as top performing in the original benchmark (Nikitin et al. 2026 Sep 17), the ComBat family methods demonstrated highest prediction quality while maintaining biomarker correlations in our current study.

We also derived specific thresholds for the correlation metrics (L and M classes) in this study. For the correlation between expression before and after harmonization, we applied a permissive threshold of 0.75 in order to cut off methods that corrupt biology (MNN and Scanorama). After that, we required cross-batch agreement biomarkers in the same biology class to increase compared to the raw expression, and cross-batch agreement of different biology to decrease – a stringent criterion that passed only 61 of the 2,150 non-raw approaches (2.8%).

Finally, we proposed a benchmarking workflow (Figure 7) that encompasses the metrics and thresholds we derived in this study. We hope the pipeline and the harmonization quality control approaches we derived here will be utilized for batch effect correction elsewhere in transcriptomics, improving batch effect correction.

## Data availability

The expression matrices, cohort annotation and harmonization metric tables analyzed here are deposited at Zenodo and accessible via the ling https://doi.org/10.5281/ZENODO.22737294. The derived tables behind the metrics in this article are provided as Supplementary File 1 and Supplementary File 2 and in the same record. Source accession numbers for the public cohorts are listed in Supplementary File 1 of the previous benchmark [38].

## Software availability

Source code to assemble the tables and conduct the analysis is deposited in GitHub in the repository https://github.com/Nikit357/FL_harmonization, folder article_2_extended_comparison/. The ComboBatch harmonization tool is available via the GitHub link https://github.com/Nikit357/ComboBatch, archived at Zenodo (https://doi.org/10.5281/ZENODO.22755758). The containerized Shambhala-2 implementation is released in GitHub at https://github.com/Nikit357/Shambhala2_fast, archived in Zenodo (https://doi.org/10.5281/ZENODO.22756647).

## Supplementary material

**Supplementary Figure 1. Biomarker panel coverage and correlation.** (A) Scatterplot of the coverage of biomarker genes over all harmonization attempts of the source benchmark (fraction of attempts in which the gene had defined expression value) against its coverage over the unharmonized gene expression matrices, color annotated by the primary cell type of the gene. The dashed line is the identity line; genes below it are lost by harmonization and not by platform. (B) Violin plots of the within-cohort Spearman ρ between harmonized and unharmonized expression of each gene over all approaches, grouped by the gene set of primary membership and ordered by median ρ; the lines inside the violins mark the median and the quartiles (Supplementary File 3, sheet Rho_by_signature).

**Supplementary Figure 2. Variance explained by the first ten principal components and its batch share.** (A) Heat map of the percentage of total variance explained by each of PC1 to PC10, averaged over the approaches of each harmonization method, with methods ordered from the highest mean PCReg RNA_BATCH (top) to the lowest. The 15 clustermap best approaches are averaged into one additional row, “clustermap best”, shown in bold. (B) Heat map of the RNA_BATCH R² of each of PC1 to PC10, in percent, over the same rows. (C) The heat map of panel A with rows by batch removal strategy. (D) The heat map of panel B with rows by batch removal strategy. Blue color scales give the percentage of variance and red color scales, from 0 to 100%, give the batch R².

**Supplementary Figure 3. Principal component regression metrics comparison.** (A) Barplot showing PCReg metrics by 3 columns visualized by harmonization methods, sorted by PCReg RNA_BATCH. Color annotations below the bars show method harshness score and overall quality. Best group means a composed group of the clustermap best approaches. (B) The analogous barplot for 4 PCReg metrics by batch removal strategies. Higher values indicate better overlap of classes by a given column in the PCA space. Error bars indicate 95% confidence interval (CI) estimation.

**Supplementary Figure 4. Comparison of kBET and LISI against PCReg.** (A) Scatterplot of LISI vs PCReg by RNA_BATCH, color annotated by harmonization methods. (B) The same scatterplot color annotated by batch removal strategies. (C) Scatterplot of LISI vs PCReg by Diagnosis_cell_type_unified, color annotated by harmonization methods. (D) The same scatterplot color annotated by batch removal strategies. (E) Scatterplot of kBET acceptance rate vs PCReg by RNA_BATCH, color annotated by harmonization methods. (F) The same scatterplot color annotated by batch removal strategies. The clustermap best approaches are shown as empty black stars on all the panels of this figure.

**Supplementary Figure 5. Additional global metrics analysis.** (A) Scatterplot showing correlation of PCReg PLATFORM_RNA with PCReg RNA_BATCH. (B) Scatterplot of PCReg COHORT_LABEL versus PCReg RNA_BATCH. (C) Scatterplot of DSC RNA_BATCH versus PCReg RNA_BATCH. (D) Scatterplot of DSC PLATFORM_RNA versus DSC RNA_BATCH. (E) Scatterplot of DSC COHORT_LABEL versus DSC RNA_BATCH. Scatterplots in panels A–E are color annotated by batch removal strategy. (F) Scatterplot of tSNE centroid dispersion RNA_BATCH versus PCReg RNA_BATCH, color annotated by batch removal strategy. (G) The same scatterplot of tSNE centroid dispersion RNA_BATCH versus PCReg RNA_BATCH, color annotated by harmonization method. (H) The same scatterplot with color annotation by post-removal and connection lines that show transition between approaches with and without post-removal. (I) Scatterplot of ASW RNA_BATCH versus PCReg RNA_BATCH, color annotated by batch removal strategy. (J) The same scatterplot of ASW RNA_BATCH versus PCReg RNA_BATCH, color annotated by harmonization method. Black empty stars on all the panels of this figure show the clustermap best approaches.

**Supplementary Figure 6. Global metrics comparison by imputation, harmonization methods and batch removal strategies.** (A) Boxplots of PCReg RNA_BATCH by harmonization methods stratified by imputation. Methods are ordered by mean PCReg RNA_BATCH across all batch removal strategies, imputations and post-removals. The clustermap best approaches are shown in bold. (B) Boxplots of PCReg RNA_BATCH by harmonization methods stratified by batch removal strategies. Harmonization methods order is the same as in panel A, but the group of clustermap best approaches is empty for strategies where no best approaches were selected. (C) Boxplots of PCReg RNA_BATCH by harmonization methods stratified by post-removal. The group of clustermap best approaches is empty in the post-removal boxplot because only the approaches without post-removal were selected. (D) Scatterplot showing correlation of PCReg RNA_BATCH and RNA_BATCH R² by first PC, color annotated by harmonization methods with black stars indicating the clustermap best approaches. (E) The same scatterplot color annotated by batch removal strategies.

**Supplementary Figure 7. LISI metrics of local neighborhood.** (A) Barplot of the LISI metrics by 3 columns (Diagnosis_cell_type_unified, COHORT_LABEL, RNA_BATCH) visualized by harmonization methods, sorted by iLISI RNA_BATCH. Color annotations below the bars show method harshness score and overall quality. Best group means a composed group of the clustermap best approaches. (B) The analogous barplot for 4 LISI metrics by batch removal strategies (added the metric for the RNASEQ_SOURCE column). Higher values indicate better overlap of classes by a given column in a local nearest neighbors’ environment. Error bars indicate 95% confidence interval (CI) estimation. (C) Heatmap of LISI by RNA_BATCH, clustered by harmonization methods (columns) and batch removal strategies (rows). Each cell shows the metric value averaged by imputations and post-removals. (D) The similar heatmap for LISI by Diagnosis_cell_type_unified. Best approaches are marked as black asterisks in the panels C and D.

**Supplementary Figure 8. Analysis of a more diverse set of local mixing metrics: LISI, kBET acceptance rate, tSNE and UMAP entropy.** (A) Scatterplot showing LISI and kBET acceptance rate by RNA_BATCH, color annotated by harmonization methods. (B) The same scatterplot color annotated by batch removal strategies. (C) Barplot of the kBET metrics by 3 columns (PLATFORM_RNA, COHORT_LABEL, RNA_BATCH) visualized by harmonization methods, sorted by kBET acceptance rate RNA_BATCH. Color annotations below the bars show method harshness score and overall quality. Best group means a composed group of the clustermap best approaches. (D) The analogous barplot for the same 3 kBET metrics by batch removal strategies. Higher values indicate better overlap of classes by a given column in a local nearest neighbors’ environment. Error bars indicate 95% confidence interval (CI) estimation. (E) Scatterplot of tSNE entropy vs LISI by RNA_BATCH, color annotated by harmonization methods. (F) The same scatterplot color annotated by batch removal strategies. (G) Scatterplot of tSNE entropy vs tSNE centroid dispersion by RNA_BATCH, color annotated by harmonization methods. (H) The same scatterplot color annotated by batch removal strategies. (I) Scatterplot of UMAP entropy vs tSNE entropy by RNA_BATCH, color annotated by harmonization methods. (J) The same scatterplot color annotated by batch removal strategies. (K) Scatterplot of UMAP centroid dispersion vs tSNE centroid dispersion by RNA_BATCH, color annotated by harmonization methods. (L) The same scatterplot color annotated by batch removal strategies. The clustermap best approaches are shown as empty black stars on the panels A, B and E – L.

**Supplementary Figure 9. Distributional and other metrics performance.** (A) Barplot of the KS fraction significant metrics by 3 columns (PLATFORM_RNA, COHORT_LABEL, RNA_BATCH) visualized by harmonization methods, sorted by KS fraction significant RNA_BATCH. Color annotations below the bars show method harshness score and overall quality. Best group means a composed group of the clustermap best approaches. (B) The analogous barplot for the same 3 KS metrics by batch removal strategies. Higher values indicate higher proportion of pairwise comparisons with significant differences in gene expression distributions by Kolmogorov-Smirnov test. Error bars indicate 95% confidence interval (CI) estimation. (C) Scatterplot of graph connectivity by Diagnosis_cell_type_unified versus PLATFORM_RNA, color annotated by harmonization methods. (D) The same scatterplot color annotated by batch removal strategies. (E) Scatterplot of distance ratio (intragroup to intergroup) by Diagnosis_cell_type_unified versus RNA_BATCH, color annotated by harmonization methods. (F) The same scatterplot color annotated by batch removal strategies. (G) Scatterplot of Watermelon score by Diagnosis_cell_type_unified versus RNA_BATCH, color annotated by harmonization methods. (H) The same scatterplot color annotated by batch removal strategies. (I) Scatterplot of mean Watermelon score by biology columns versus the same score by batch columns, color annotated by harmonization methods. (J) The same scatterplot color annotated by batch removal strategies. The clustermap best approaches are shown as empty black stars on the panels C – J. (K) Heatmap showing gene expression quantiles and standard deviation by batch removal strategies.

**Supplementary Figure 10. Barplots comparing metrics of different harshness harmonization methods by batch removal strategies.** (A) PCReg RNA_BATCH. (B) PCReg Diagnosis_cell_type_unified. (C) tSNE entropy RNA_BATCH. (D) LISI RNA_BATCH. (E) LISI Diagnosis_cell_type_unified. (F) KS fraction significant RNA_BATCH. In each barplot, the harmonization strategies are sorted by the metric shown, and the clustermap best approaches are shown as a surrogate batch removal strategy.

**Supplementary Figure 11. Stacked barplots showing impact of metric type, group and column on the equally weighted composite score.** (A) Metric type by batch removal strategy. (B) Metric type by harmonization method. (C) Metric group by batch removal strategy. (D) Metric group by harmonization method. (E) Metric column by batch removal strategy. (F) Metric column by harmonization method. In all barplots bars are sorted by average value descending.

**Supplementary Figure 12.** Individual approaches sorted by the composite score, with color bars for the strategy, the imputation, the method, the post-removal setting and the method harshness tier of each approach. The clustermap best approaches are shown with black stars.

**Supplementary Figure 13. Biology and prediction metric classes L, M and N across the three method harshness tiers.** Each panel shows one metric as boxplots over the approaches by harshness tiers. The p value on each panel is the raw p value (not corrected for multiple testing) of the Kruskal–Wallis H test across the three tiers, computed from the chi-square approximation of H with two degrees of freedom. Number of approaches per tier for every metric is given in Supplementary File 1, sheet Harshness_full. (A) Class L mean within-cohort marker correlation before and after harmonization. (B) Class L marker correlation minus the housekeeping-gene correlation. (C) Class M same-biology cross-batch biomarker rank agreement. (D) Class M different-biology cross-batch rank agreement. (E) Class M margin, same-biology minus different-biology agreement. (F) Class M margin corrected by the unharmonized baseline at the same strategy and imputation. (G) Class N LOBO three-class macro F1 over all folds. (H) The same metric over the multiclass folds only. (I) Class N LOBO three-class macro AUC, which is defined on multiclass folds only. (J) Class N LOBO two-class macro F1 over all folds. (K) The same metric over the multiclass folds only. (L) Class N LOBO two-class macro AUC, defined on multiclass folds only. In panels G, I, J and L, the gray box beside each tier is the label-permutation background: for each approach, the mean of the same metric over 100 permutations of the biology labels. Boxes span the IQR with the median marked by a horizontal line; whiskers extend to 1.5 × IQR and points beyond them are drawn individually.

**Supplementary Figure 14. Relationships between the metrics of classes L, M and N.** Each row is two scatterplots for one pair of metrics, color annotated by harmonization method (left column: A, C, E) and by batch removal strategy (right column: B, D, F). (A, B) LOBO three-class macro F1 over all folds against the class L mean marker correlation. (C, D) LOBO three-class macro F1 over all folds against the class M margin. (E, F) LOBO three-class macro F1 over the multiclass folds only against the class M margin gain over the unharmonized baseline. The ρ and p above each panel are the Spearman correlation and its raw two-sided p over all approaches that contain both metrics. The clustermap best approaches are shown as empty black stars on all panels. Spearman correlations for all 55 pairs of the L, M and N metrics are given in Supplementary File 1, sheet Scatter_correlations.

**Supplementary Figure 15. Cross-correlation of the harmonization quality metrics.** Clustered heat map of the pairwise Spearman correlation between 325 metrics, computed over the 2,234 approaches with pairwise-complete observations. Rows and columns are ordered by average-linkage hierarchical clustering, with the distance between two metrics taken as one minus the Pearson correlation of their rows. The three color bars beside the rows and above the columns show, from the outer to the inner bar, the metric group of the registry, the metric type (global, local, distributional similarity or other) and the detailed metric class (global, local, distributional, structural, L, M, N or unassigned); the assignment of every metric is listed in Supplementary File 1, sheet Metric_class_assignment.

**Supplementary Figure 16. Harmonized against unharmonized expression for four linear and four non-linear harmonization methods.** Each row is one method under the no-removal strategy (S0_no_removal) with KNN imputation and without post-removal: (A) 03_limma, (B) 05_combat, (C) 29_combat_ref and (D) 13_fsmvn, linear methods, and (E) 16_fsqn_r, (F) 17_quantile, (G) 10_mnn and (H) 12_scanorama, non-linear methods. The columns show the three genes: BCL6, a GC B-cell marker, IRF4, a post-GC marker, and EEF1A1, a housekeeping gene. Each point is one sample of the four largest cohorts (PUB_DLBCL_GSE117556, SOM, PUB_DLBCL_NCICCR and PUB_DLBCL_GOYA), color annotated by cohort, with the unharmonized log2 expression (01_raw at the same strategy and imputation) on the x axis and the harmonized value on the y axis. The dashed line is the identity line. The ρ in each cell is the median over the four cohorts of the within-cohort Spearman correlation between harmonized and unharmonized values, the quantity that class L averages over the biomarker panel.

**Supplementary File 1. The analysis set, its metric classes and the statistical tests.** Analysis_set: the 2,234 approaches, one row per approach, with the key columns, the flags used in the text and the metric values. Canonical_set_reconciliation: the reconciliation from the 2,407 completed approaches of the source benchmark (Nikitin et al. 2026 Sep 17) to 2,234. Metric_class_assignment: the 344 metrics of the registry, by the name they carry in Supplementary File 2 of the source benchmark (which holds their polarity, group and type), with a scoring and a metadata flag, their availability in the pinned snapshot and their Article 2 metric class. Class_percentile_scores: for each of the 2,234 approaches, the score of every metric class (the mean rank percentile of its metrics) and whether the class elected it. Generalizability_ranking: the 2,234 approaches ordered by the generalizability index, with its six components. Method_census_by_class: the number of elected places that each harmonization method takes in each metric class. Strategy_and_imputation_summary: the metric medians grouped by batch removal strategy and imputation.

Harshness_by_LMN: the Kruskal–Wallis tests of the class L, M and N metrics across the three harshness tiers. Harshness_full: the same test for every numeric class L, M and N column, 186 metrics, with the mean, median and median absolute deviation (MAD) and the number of approaches per tier and the pairwise Mann–Whitney p values. Paired_tests: the 285 hypothesis tests behind the comparisons in the text, each with its groups, group sizes, medians, statistic and raw p value. Scatter_correlations: the 97 Spearman correlations for the scatterplot relationships, with n, ρ and p. Composite_dispersion: the contribution of each metric type and metric group to the composite score by strategy and by method, with the range and the standard deviation of the group means. PCA_component_means: the mean percentage of total variance and the mean RNA_BATCH R^2^ on each of the first ten principal components, for every method, every strategy, the clustermap group and the unharmonized no-removal approaches.

**Supplementary File 2. Cross-batch structure, the best approach sets and prediction.** Cross_election_matrix: the Jaccard index between the elected sets of every pair of the nine sets (eight metric classes and the 15 clustermap best approaches). Cross_election_pairs: the same 36 pairs with the set sizes, the number of shared approaches, the union, the Jaccard index, the overlap expected by random background and the hypergeometric p values for more and for less overlap than chance. Elected_sets_by_class: the approaches elected by each set. Agreement_specific_61: the 61 approaches that raise same-biology agreement and lower different-biology agreement against their unharmonized baseline, with their class L, M and N values. Margin_raw_and_delta: the class M columns of all 2,234 approaches, in raw form and with the unharmonized baseline subtracted. LOBO_summary: the class N metrics on both fold cuts and for both targets (2 and 3 classes) for the selected groups of approaches and for the leading approaches. Per_batch_folds: the macro F1 and AUC of every held-out batch for the prediction best and the clustermap best approaches, for both targets. Fold_composition: for each held-out batch and number of biology classes, the number of folds and their median F1, AUC and sample count, pooled over approaches. Threshold_sensitivity: the overlap of every pair of elected sets when the election takes the top 2.5%, 5% or 10% of approaches, with the Jaccard index. It also gives the rank of each Jaccard index among the 36 pairs and the Spearman correlation of each threshold’s Jaccard values with those at 5%.

**Supplementary File 3. The biomarker gene panel detailed tables.** Panel_coverage_by_attempt: the number of biomarker genes retained as the coverage threshold is changed. QC_gate_sweep: the number of marker genes that pass the per-gene quality gate over the grid of the two thresholds. Rho_by_signature: the per-gene summary of the within-cohort Spearman ρ with the signature membership and the saturation flag of each gene, for the biomarker genes measured in at least one approach. Narrow_set_56_genes: the genes of the 56-gene narrow panel with their annotation.

## Ethics and consent

Ethical approval and consent were not required. The study used de-identified transcriptomic profiles that had already been published and produced no new human material or data.

## Author contributions

Daniil Nikitin: conceptualization, methodology, software, formal analysis, investigation, data curation, validation, visualization, writing – original draft, writing – review and editing. Arsen Arakelyan: conceptualization, methodology, supervision, project administration, funding acquisition, writing – review and editing.

## Acknowledgements

We thank Alisa Sadekova and Alexander Ryabykh for help with dataset assembly. For discussion we thank Andrey Kravets, Anastasiya Yudina, Daniil Ivanov, Konstantin Chernyshov, Aleksei Efremov and Katerina Nuzhdina.

